# Predicting the translation efficiency of messenger RNA in mammalian cells

**DOI:** 10.1101/2024.08.11.607362

**Authors:** Dinghai Zheng, Logan Persyn, Jun Wang, Yue Liu, Fernando Ulloa Montoya, Can Cenik, Vikram Agarwal

**Author notes:** These authors contributed equally to this work (Dinghai Zheng, Logan Persyn, Jun Wang). Correspondence to Can Cenik and Vikram Agarwal.

## Abstract

The degree to which translational control is specified by mRNA sequence is poorly understood in mammalian cells. Here, we constructed and leveraged a compendium of 3,819 ribosomal profiling datasets, distilling them into a transcriptome-wide atlas of translation efficiency (TE) measurements encompassing >140 human and mouse cell types. We subsequently developed RiboNN, a multitask deep convolutional neural network, and classic machine learning models to predict TEs in hundreds of cell types from sequence-encoded mRNA features, achieving state-of-the-art performance (r=0.79 in human and r=0.78 in mouse for mean TE across cell types). While the majority of earlier models solely considered 5′ UTR sequence^1^, RiboNN integrates contributions from the full-length mRNA sequence, learning that the 5′ UTR, CDS, and 3′ UTR respectively possess ∼67%, 31%, and 2% per-nucleotide information density in the specification of mammalian TEs. Interpretation of RiboNN revealed that the spatial positioning of low-level di- and tri-nucleotide features (*i.e.*, including codons) largely explain model performance, capturing mechanistic principles such as how ribosomal processivity and tRNA abundance control translational output. RiboNN is predictive of the translational behavior of base-modified therapeutic RNA, and can explain evolutionary selection pressures in human 5′ UTRs. Finally, it detects a common language governing mRNA regulatory control and highlights the interconnectedness of mRNA translation, stability, and localization in mammalian organisms.

## INTRODUCTION

Protein abundances are determined by the complex interplay of steady-state mRNA levels, mRNA translation rates, and protein turnover rates. Numerous machine learning models have been developed to model the sequence-encoded features that influence steady-state levels of mammalian mRNAs from both the perspectives of transcriptional regulation^2–6^ and mRNA turnover^7^. However, most attempts to model translational regulation from mRNA sequence have focused on bacteria and yeast^8–13^. Although such models do exist for mammals, most focus on the functional roles of specific regions such as the 5′ untranslated region (5′ UTR)^14–16^ or coding region sequence (CDS)^17–19^, despite the recognition that the full mRNA sequence (*i.e.*, including 3′ UTRs) jointly influences translation^1,20,21^. Several models consider full-length mRNA, but have either only implicitly modeled translational regulation^22,23^, or have evaluated only a limited set of cell types while achieving modest performance (r^2^≈0.40)^24,25^. Modeling translational regulation more precisely among diverse cell types would elucidate the functional consequences of synonymous, missense, and non-coding mutations in mRNA. Consequently, this would advance the goals of identifying the mechanistic underpinnings of ribosome occupancy and protein abundance quantitative trait loci (rQTL and pQTL, respectively)^26,27^, diagnosing pathogenic genetic variants, and designing more translationally competent mRNA therapeutics and gene therapies.

Global translation rates can be estimated through several strategies, including: i) fitting translation rate parameters from differential equations, using measurements of mRNA and protein abundances as well as mRNA half-life^28,29^; ii) computing protein-to-mRNA ratios (PTRs)^22,23,30^; iii) polysome profiling, in which ribosomal fractions are run on a sucrose gradient and mRNAs within each fraction are sequenced to estimate their approximate ribosomal loading^14,15,21,31^; and iv) ribosome profiling (*i.e.*, Ribo-seq), normalizing ribosome density to RNA abundance as a metric for TE^32^. Of these techniques, the first two strategies are both indirect estimates of translation rate. Importantly, inferred translation rates from the differential equation modeling strategy were shown to be poorly related to experimentally measured rates^33^, limiting the accuracy of this approach. Moreover, PTRs are partially confounded by protein degradation rates and protein secretion^22,23,30^. Therefore, of these four methods, polysome and ribosome profiling are considered more direct methods of assessing translation rates^33^.

In eukaryotes, translation is regulated at the initiation and elongation steps^34,35^, which can be modulated by *cis*-acting sequences. In particular, *cis*-regulation of translation initiation has historically been the focus due to its recognition as the rate-limiting step of translation^36^. The propensity for secondary structure near the 5′ mRNA cap, the sequence context of the translation initiation codon, presence of upstream short open reading frames (ORFs), and binding sites for various RNA-binding proteins provide concrete mechanisms of translational regulation via *cis*-acting elements predominantly in 5′ UTRs^37^. Importantly, the protein coding sequence is also a key determinant of TE. Relatively more is known in unicellular organisms; in particular, codon usage differs significantly across genes, with more abundant proteins utilizing a biased set of codons^38,39^. The most widely recognized mechanism for codon-specific influence on translation relates to differences in the active pool of corresponding tRNAs^40–42^. Coding sequence differences are also suggested to impact protein expression through secondary structure-mediated mechanisms that do not correlate with tRNA abundance^43^. Moreover, non-synonymous coding variants can alter translation independently from tRNA abundance, translation initiation efficiency, or overall mRNA structure via the interaction of the encoded peptide with the ribosome exit tunnel^44^. Parallel work in vertebrate organisms established a link between translation and RNA stability; for instance, certain codons that slow down translation are associated with unstable mRNA^17,45–50^. Taken together, these studies reveal that the entire mRNA sequence can potentially modulate translation through a variety of mechanisms. However, the contribution of specific functional regions in determining translation of endogenous mRNAs has yet to be described quantitatively. A precise measurement of translation rate would enable a clear-eyed examination of how different sequence properties and functional regions modulate translation rates relative to one another.

Despite the widespread abundance of ribosomal profiling datasets, attempts to examine the relative contribution of sequence and structural features to the specification of translation rate have been hampered by their inaccessibility in a unified resource. In this study, we systematically assembled a compendium of 1,282 human and 995 mouse ribosome profiling datasets, matched to corresponding RNA-seq data, to derive more precise TE measurements in mammalian cells. This effort reflects the synthesis of the largest and most comprehensive compendium of TE measurements ever assembled to date. Using enhanced measurements of TE, we derived improved sequence-based models towards the goal of improving the predictability of TE from RNA sequence. Our state-of-the-art model RiboNN, a deep convolutional neural network, is capable of predicting the effects of RNA sequences (*e.g.*, including base-modified, therapeutically delivered mRNA) on translational regulation, in agreement with functional measurements derived from massively parallel reporter assays and population genetic data demarcating regions of evolutionary constraint. RiboNN reconciles several limitations of existing models, possessing the following properties: i) it models the impact of the full-length mRNA sequence on TE in numerous cell types, ii) it exhibits superior performance in predicting TE from mRNA sequence, iii) it identifies the location-dependent effects of short, di- and tri-nucleotide features (*i.e.*, including codons) as the key sequence features explaining model performance, and iv) it helps to quantify the relative contributions of different functional regions on TE, a feat which has largely been evaluated qualitatively in the past. Finally, it postulates the existence of a common language underpinning mRNA translation, stability, and localization in mammalian organisms.

## RESULTS

### Preparation of a compendium of human and mouse TE datasets from ribosome profiling data

To construct a comprehensive, high-quality dataset of TE measurements, we systematically compiled 3,819 human and mouse ribosome profiling datasets from the GEO database. We filtered these into 1,282 human and 995 mouse samples representing matched ribosome profiling and RNA-seq data from numerous tissues and cell types. We then uniformly processed the datasets using an open-source bioinformatics pipeline^51^. We required each sample to pass the following quality control filters: i) ≥70% of ribosome-protected fragments (RPFs) mapped to the CDS, and ii) transcripts globally had a minimum average read coverage of 0.1x (detailed in companion manuscript^52^). This yielded 1,076 human and 835 mouse ribosome profiling datasets. We then calculated TE using a compositional regression approach that overcomes the mathematical biases associated with the commonly used log-ratio approach^53,117^ (**Fig. 1a; Methods**). We summarized the datasets by averaging TEs across samples belonging to the same cell types, yielding matrices of 10,348 genes x 78 cell types for the human and 10,870 genes x 68 cell types for the mouse (**Fig. 1a**, **Supplementary Table 1**). Each cell type varied in quality with respect to the number of missing genes (**Supplementary Fig. 1**), in part due to factors such as variable sequencing depth and number of samples averaged. This resource enabled us to assess the degree to which TEs are similar among different mRNAs across cell types. We calculated the Spearman’s correlation coefficient (rho) between the TEs of transcripts across all possible pairs of human cell types (**Fig. 1b**). We observed that most of the cell types were highly correlated to each other, with a small subset possessing low correlation to most other cell types (**Fig. 1b**). This subset appeared to have lower data quality, as measured by a low median read coverage, leading to a large proportion of missing values (**Fig. 1b**). The high correlation between most cell types is suggestive of common translational regulation mechanisms across most cell types. Parallel results were observed for the inter-cell-type comparisons in the mouse (**Extended Data Fig. 1a**).

**Fig. 1.**
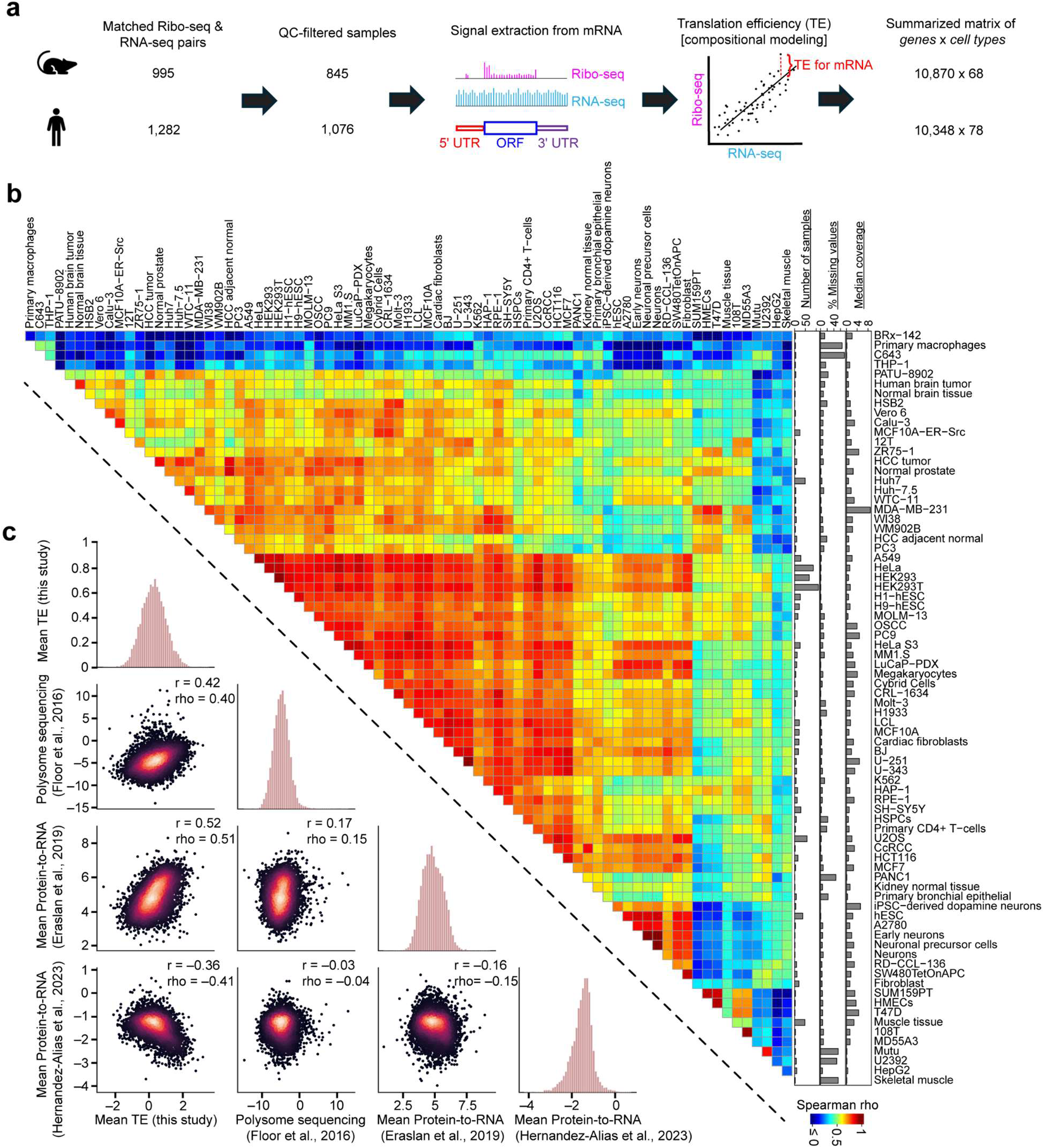
Integrative analysis of thousands of human and mouse ribosomal profiling datasets measuring TE. **a,** Schematic showing the workflow of transcriptome-wide TE calculations for the human and mouse, using paired RNA-seq and ribosome profiling datasets. **b,** Heatmap of Spearman correlation coefficients comparing TEs derived from each pair of 78 human cell types. Cell types are clustered using hierarchical clustering. Right panel barplots show quality control data for the human cell type shown in each row. **c,** Comparison of mean TEs (*i.e.*, averaged across human cell types) for mRNAs derived from this study relative to alternative measurements of translational output measured in prior studies^21,23,30^. The Pearson (r) and Spearman (rho) correlation coefficients between each pair of measurements is also shown.

To validate the biological relevance of TEs relative to other methods to measure translational regulation, we compared the TE across cell types with previously reported PTR ratios^23,30,54^ and ribosome load (number of ribosomes per transcript), as measured by polysome sequencing in HEK293T cells^21^. We normalized the ribosome load to CDS length because longer CDSs can accommodate more translating ribosomes. Given the strong correlation based upon dataset of origin (**Supplementary Fig. 2**), we evaluated the relationship between the means of each dataset. The ribosome load and mean PTR across tissues^23^ were positively correlated with our mean TE (r=0.42, rho=0.4 and r=0.52, rho=0.51, respectively; **Fig. 1c**). However, the mean PTR reported from a recent study^30^ was weakly negatively correlated with our mean TE (r=−0.36, rho=−0.41; **Fig. 1c**). These PTR measurements were highly discordant with other datasets as well, suggesting that the most parsimonious explanation to be the relatively lower reliability of this PTR dataset^30^. Even stronger correlations were observed between mouse mean TE and ribosome load in mouse 3T3 cells^31^ (r=0.61, rho=0.64; **Extended Data Fig. 1b**). Together, these results suggest that our TE scores are informative of protein synthesis rates in both organisms.

### Classical machine learning models to predict TE

To evaluate the predictability of our TE measurements, we trained regression models on pre-computed sets of sequence-encoded features derived from the mRNA. The feature sets considered include: i) the lengths of the 5′ UTR, CDS, 3′ UTR, and entire transcript; ii) nucleotide frequencies of all regions; iii) codon frequencies; iv) amino acid frequencies; v) k-mer frequencies of length 2 to 6 in the 5′ UTR, CDS, and 3′ UTR regions; vi) the frequency of each nucleotide found in the wobble position; vii) the nucleotide identity at the −3, −2, −1, +4, and +5 Kozak positions; viii) dicodon counts found to affect TE in yeast^42^; and ix) multiple secondary structure features. For benchmarking purposes, we also considered biochemical features, defined as those derived from experimental measurements such as CLIP-seq and RIP-seq^7^ (**Methods**).

To identify which feature sets usefully contributed to prediction of mean TE across all human cell types, we used an iterative method that compared the cross-validated (CV) performance of a light gradient-boosting machine (LGBM) model trained with a specific feature set to one trained without it. If the model including the feature set performed statistically significantly better on ten held-out data folds than the model without it, that feature set was deemed useful (**Methods**). The feature sets found to be useful include: i) regional and total sequence lengths; ii) UTR nucleotide frequencies; iii) codon frequencies; iv) amino acid frequencies; and v) the 3-mer frequencies of the 5′ UTR (**Fig. 2a**). All remaining feature sets did not further contribute to TE prediction (“Other” in **Fig. 2a**), including secondary structure features, in contrast to prior findings^43^.

**Fig. 2.**
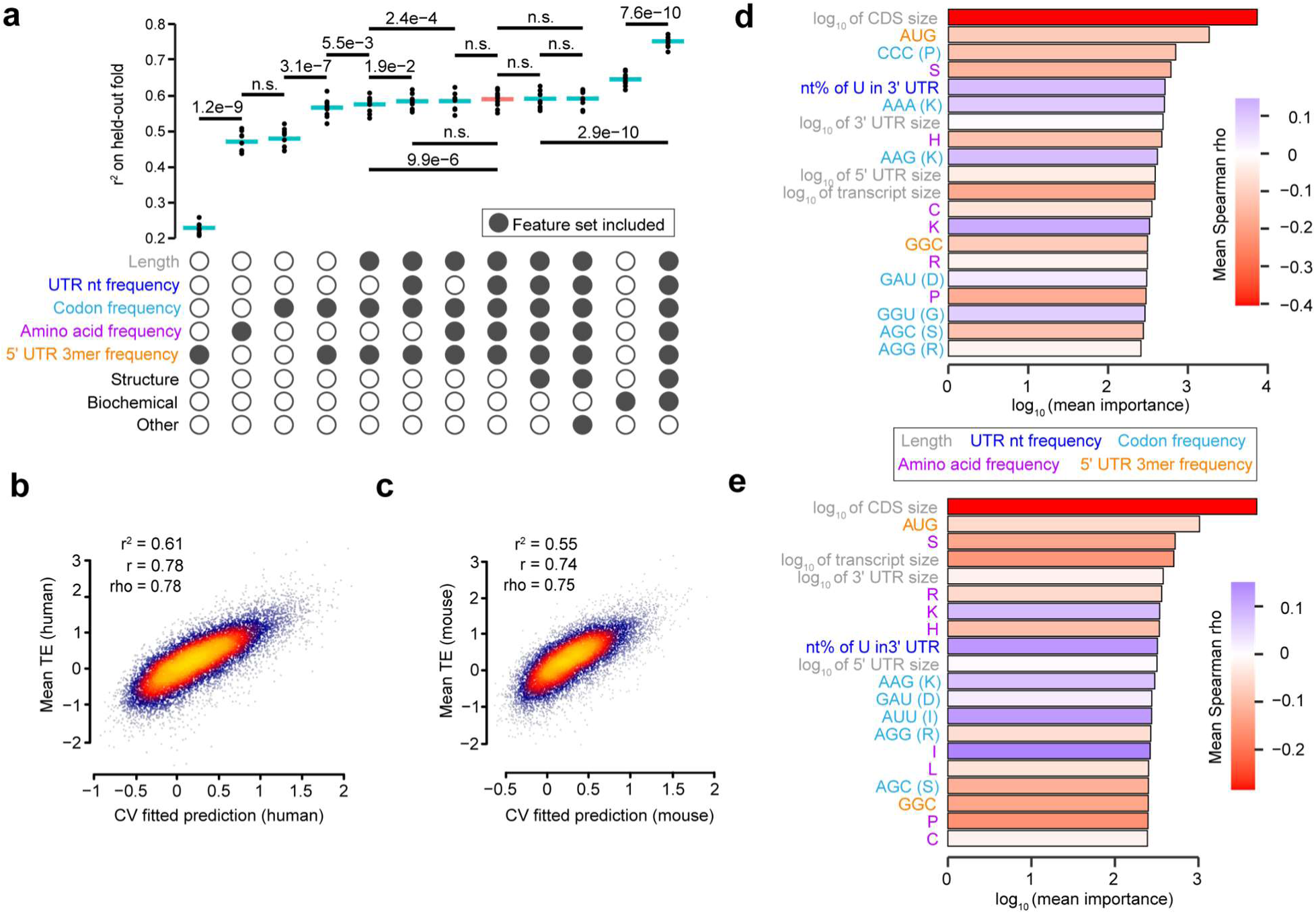
A classical machine learning approach to predict mammalian TEs from mRNA sequence. **a,** UpSet plot showing the r^2^ measured on ten held-out CV folds of LGBM models which predict the mean TE across human cell types using various feature sets. Colored feature sets are indicative of those that contributed to the optimal sequence-only model. Median r^2^ and statistically significant differences in performance between pairs of models are indicated. P-values were calculated using one-sided, paired t-tests adjusted with a Bonferroni correction. All additional feature sets considered, but that did not have a significant improvement on performance, are labeled as “Other”. **b-c,** Scatter plots comparing the predicted and observed mean TEs, averaged across cell types, for both the human (**b**) and mouse (**c**). The r^2^, Pearson (r), and Spearman (rho) correlation coefficients, integrating the results across ten CV folds, are also shown. **d-e,** Importance of the features used by the optimal sequence-only model (shown as a red bar in panel **(a)** for both the human **(d)** and mouse **(e)**. For a given feature, importance was measured as the sum total information gain across all splits using the feature, averaged across all folds. The colors of the bars correspond to the mean Spearman rho, averaging rho values between the features and TE values from each cell type. Feature names are colored according to the feature set to which they belong.

Given this set of selected features, we compared three additional machine learning approaches to assess their relative performance: lasso, elastic net, and random forest. We confirmed that LGBM performed the best (**Supplementary Fig. 3a**). We then trained LGBM models on all 78 human and 68 mouse cell types. The correlation between the mean TE and average over the predictions of each cell type was r=0.78 for human and r=0.74 for mouse (**Fig. 2b-c**). The r^2^ (averaged across the held-out folds) for predicting the mean TE across cell types was 0.61 and 0.55 for the human and mouse, respectively (**Fig. 2b-c**). Cell types with poorer data quality, such as a lower fraction of detectable genes, generally led to models with inferior performance (**Supplementary Fig. 4**). Although the hand-crafted feature sets could not easily include positional information, the regression models were still able to achieve impressive performance. We benchmarked our human LGBM model against two prior models, both of which consider only 5′ UTR information and were trained on HEK293T data. We found that Optimus^14^ achieved 4−6-fold inferior results (r^2^=0.12 on HEK293T TE; r^2^=0.10 on mean TE) relative to LGBM; similarly, the FramePool^16^ model also achieved 2−4-fold inferior results (r^2^=0.18 on HEK293T TE; r^2^=0.17 on mean TE) (**Supplementary Fig. 3b-e**).

Next, we sought to identify the relative importance of individual features for our optimal LGBM model. Several of the top-ranked features were consistent with those reported in the literature (**Fig. 2d-e**). For instance, both the human and mouse models capture: i) the known negative correlation between TE and both total mRNA sequence length and CDS length^22,55–58^, detected in polysome sequencing and PTR data as well (**Supplementary Fig. 5a**); ii) the known importance of AUG [often associated with upstream ORFs (uORFs)] and GGC trinucleotides in the 5′ UTR^59–61^; and iii) to the best of our knowledge, the previously unknown positive correlations of codons AAG (lysine) and GAU (aspartic acid) as well as the negative correlations of codons AGG (arginine) and AGC (serine). Overall, the mean feature importance across cell lines was highly correlated between the human and mouse (r=0.95, rho=0.95; **Supplementary Fig. 5b**). Taken together, these results demonstrate the robust predictive power of specific sequence-encoded features on mammalian TE, underscoring the influence of nucleotide composition and sequence length across different cell types.

### A deep neural network to predict TE from mRNA sequence

Given that deep-learning-based approaches can capture positionally aware contributions of sequence features and reveal degenerate motifs which are arduous to consider in classical machine learning models, we compared the performance of deep-learning models on the aforementioned tasks. Specifically, we trained multitask, deep convolutional neural networks to simultaneously predict TEs in all cell types examined. The input to our models consisted of a one-hot encoding of the mRNA sequence (up to a maximum of 13,318 nt), along with binary variables indicating the first reading frame of a codon for each nucleotide; the output layer consisted of multitask predictions for the TEs of either 78 human or 68 mouse cell types (**Fig. 3a**).

**Fig. 3.**
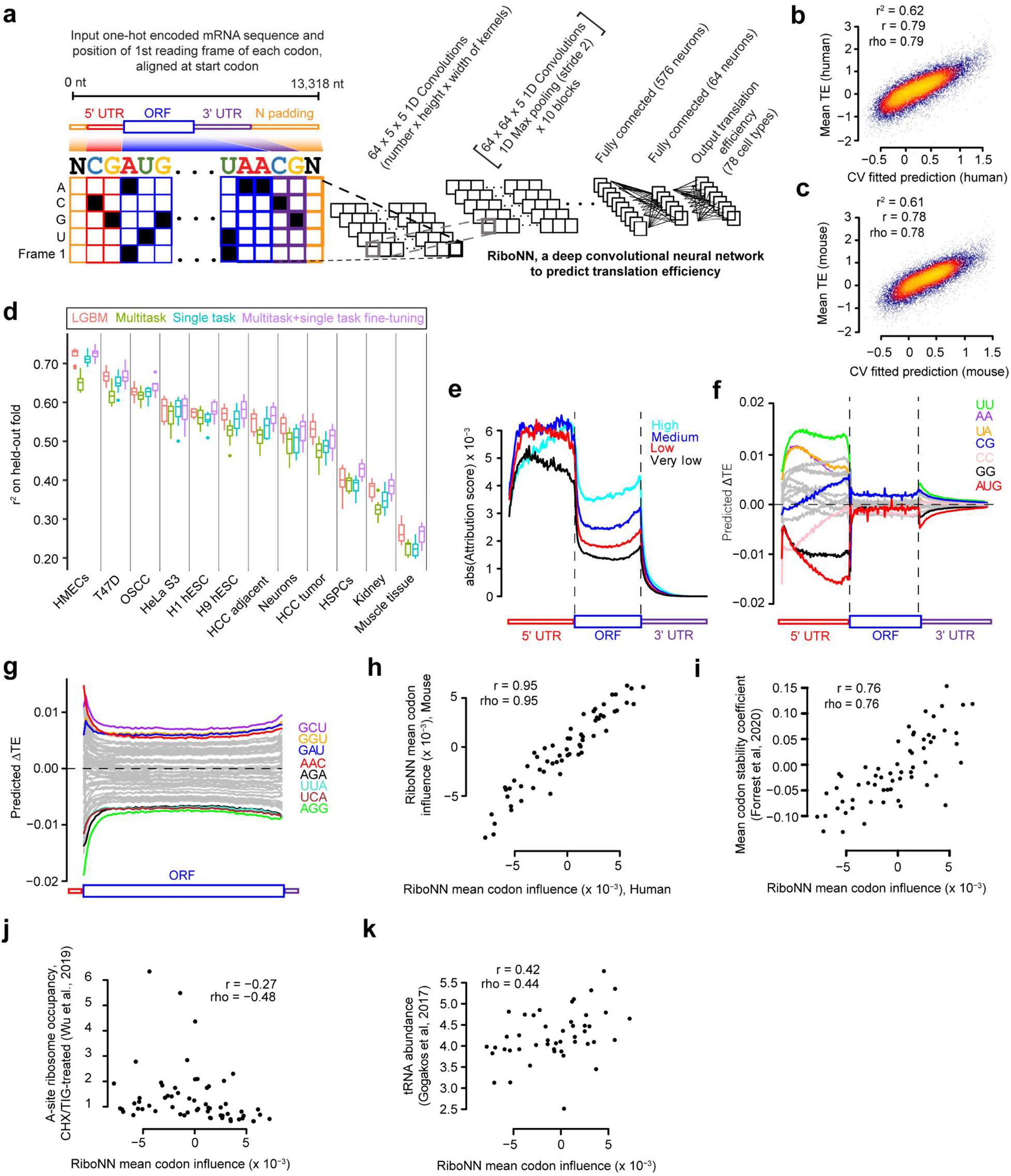
Performance and interpretation of deep learning models predicting mammalian TEs from mRNA sequence. **a,** Architecture of RiboNN, a deep multitask convolutional neural network trained to predict TEs of mRNAs in numerous cell types from an input of the mRNA sequence and an encoding of the first frame of each codon. **b-c,** Performance of RiboNN in predicting human (**b**) and mouse (**c**) mean TEs, averaged across cell types. The r^2^, Pearson (r), and Spearman (rho) correlation coefficients, integrating the results across ten CV folds, are also shown. **d,** Comparison of different model training strategies for predicting TEs in individual cell types. The following approaches were examined: LGBM trained on a single task, RiboNN trained in either a multitask or single task setting, and RiboNN trained in a multitask setting but then fine-tuned on a single task (*i.e.*, a transfer learning approach). **e,** Metagene plot summarizing the absolute value of attribution scores, averaging across all mRNAs, for percentiles along the 5′ UTR, CDS, and 3′ UTR. mRNAs were grouped into one of 4 equally sized bins according to their mean TE. **f,** Insertional analysis of 16 dinucleotides and the AUG motif. Motifs were inserted into each of 100 equally spaced positional bins along the 5′ UTR, CDS, and 3′ UTRs of each mRNA. Indicated is the average predicted change in TE for each bin plotted along a metagene. **g,** This panel is the same as panel (**f**), except it performs analysis for 61 codons (excluding the 3 stop codons) inserted into the first reading frame along the length of the CDS. **h-k,** Scatter plots showing the relationship between the codon influence (*i.e.*, the predicted effect size of each inserted codon, averaged across all positional bins) from the human RiboNN model with that of the mouse model (**h**), mean codon stability coefficients^47^ (**i**), A-site ribosome occupancy scores^62^ (**j**), and tRNA abundances^63^ (**k**). Pearson (r) and Spearman (rho) correlation coefficients are also shown.

We first repurposed a hybrid convolutional and recurrent deep neural network architecture (Saluki) designed to predict mRNA stability^7^, removing the splice site channel. In addition, we trained a new model named RiboNN, in which we removed the gated recurrent unit layer in Saluki but increased the number of convolution/max-pooling blocks from 6 to 10 to further compress mRNA sequence length by ∼1000-fold (**Fig. 3a**, **Extended Data Fig. 2**). A hyperparameter search testing a different number of such blocks and different batch sizes showed no statistical differences in performance relative to the hyperparameters we selected (**Supplementary Fig. 6**). To facilitate the learning of important features (*e.g.*, Kozak sequence) near the start codon, we fixed the start codon position in the input by aligning the mRNA sequences at the start codon. To accommodate the variability in mRNA sequence length, both the 5′ and 3′ ends of mRNAs shorter than 13,318 nt were padded with Ns (**Fig. 3a**). RiboNN achieved an r^2^ (averaged across held-out folds) of 0.62 for predicting the mean TE across the human cell types. As observed previously for LGBM models, the r^2^ degraded for cell types with poorer data quality (**Extended Data Fig. 3**). Sequence homology among intraspecies paralogs did not drastically inflate the results, because removing mRNAs in the test which were highly homologous sequences to those in the training set led to highly similar r^2^ values (**Supplementary Fig. 7**). The performance of the modified Saluki and RiboNN models were similar across cell types, with RiboNN slightly outperforming the modified Saluki (p=2.9e−10, paired Wilcoxon signed-rank test; **Extended Data Fig. 3**). Moreover, deleting the codon labels or fixing the mRNA sequences at the 5′ end (*i.e.*, rather than the start codon) each resulted in significantly lower r^2^ in most cell types (p<2.2e−16 for both paired Wilcoxon signed-rank tests; **Extended Data Fig. 3**).

We independently trained RiboNN to predict TEs in 68 mouse cell types. Like the human models, the mouse model exhibited variable performance among cell types, in a manner dependent on data quality. Overall, RiboNN achieved an r^2^ (averaged across held-out folds) of 0.61 for predicting the mean TE across mouse cell types (**Extended Data Fig. 4a**). The mouse and human RiboNN models worked almost as well when generating predictions across species as within species, suggesting an evolutionary conservation of the principles learned (**Extended Data Fig. 4b-c**). This performance could not be explained merely due to the interspecies training on orthologous sequences, because the sequence homology between mRNA pairs across species was typically weak (*i.e.*, <50%), and the variance in prediction errors for mRNAs with the highest homology was akin to those with low homology (**Extended Data Fig. 4d-e**). The final human and mouse models displayed an r^2^ of 0.62 and 0.61, respectively, in predicting mean TEs averaged across cell types (**Fig. 3b-c**), suggesting that RiboNN learned principles of translational regulation for endogenous mRNAs. Reinforcing our prior results (**Fig. 2a**), considering a set of RNA structural motifs alongside our human RiboNN model did not significantly enhance its performance relative to RiboNN alone (**Supplementary Fig. 8a-b**). However, a weak but detectable signal was apparent for RNA G-quadruplexes (RG4s) as a structural motif that could further explain the data (**Supplementary Fig. 8c**), opening the possibility that RiboNN may not comprehensively capture the effect of all such motifs.

The availability of TEs measured in various cell types provided the possibility of testing multiple modeling strategies to improve TE prediction for specific cell types. To further improve model performance, we compared single-task models and multitask models fine-tuned to a single task (*i.e.*, a transfer learning approach) on 12 randomly selected cell types exhibiting a wide distribution of r^2^ values (**Supplementary Table 2**). Interestingly, single-task RiboNN models outperformed the multitask model for most of the cell types, but were in turn outperformed by multitask models fine-tuned to a single task (**Fig. 3d**). These results highlight the power of transfer learning as an effective strategy to enable information sharing between models. Although RiboNN and LGBM displayed comparable prediction performance, RiboNN nevertheless has distinct advantages with respect to its convenient application for transcriptome-wide TE prediction, circumventing the need to pre-compute features and enabling a more computationally efficient path towards the inference of genetic variant effects. Furthermore, evaluating the features that contribute to RiboNN’s success in predicting TE may uncover novel principles of translational control that may have otherwise been overlooked.

To interpret the principles learned by RiboNN, we tested its predictive behavior in different contexts. Saliency maps are commonly utilized to explain deep learning model predictions by highlighting the input variables that contribute most towards the predicted label^64,65^. First, for each nucleotide of every human mRNA, we calculated attribution scores contributing to the prediction of mean TE across all the cell types, multiplying these with the one-hot encoding of each mRNA sequence to evaluate the predicted contribution of the input nucleotides. Averaging across all mRNAs, we generated a metagene plot using these scores, evaluating the attributed effect size (*i.e.*, absolute value) of each position along the length of each functional region of mRNA (**Fig. 3e, Extended Data Fig. 5a**). mRNAs were grouped into one of four equally sized bins according to their measured mean TE (High, Medium, Low, and Very low). This analysis revealed that 5′ UTR sequences and CDS incorporate the greatest per-nucleotide information density (∼67% and 31%, respectively) in predicting translational output, followed by the 3′ UTR having the least contribution (2%). Taking into consideration the average length of each functional region, our model predicted a total global contribution of 22%, 73%, and 5% for the 5′ UTR, CDS, and 3′ UTR, respectively. In addition, RiboNN learned position-specific contributions to TE prediction. Specifically, the identity of the first 10 codons demonstrated a ∼2-fold greater impact compared to codons positioned towards the middle of the ORF (amino acids 70 to 80) in both human and mouse (**Extended Data Fig. 5a**). These general observations were consistent for the mouse, which exhibited a 67%, 31%, and 2% per-nucleotide information density and 23%, 73%, and 4% total global contribution for the 5′ UTR, CDS, and 3′ UTR, respectively (**Extended Data Fig. 5b**). The positional importance of the early coding region was similarly greater in mice (**Extended Data Fig. 5c**), suggestive of an evolutionarily conserved principle among mammalian species.

We further examined our attribution scores using TF-MoDISco-lite^66^ to identify the most significant motifs associated with TE prediction for both human and mouse RiboNN models. Our analysis revealed that short, degenerate motifs; including CC, GG, CG, and AUGs upstream and downstream of the main ORF; are predictive of translation output (**Extended Data Fig. 5d-e**). Inspired by this finding, we performed an insertional analysis of all 16 dinucleotides and AUG to evaluate the model’s behavior upon inserting each of these short motifs along the full length of each mRNA. We observed varying influences on TE among different motifs and across different functional regions of mRNA for the same motif. Insertion of AUG and GG in the 5′ UTR demonstrated the strongest negative effect on TE prediction for both human and mouse models, while UU, AA, and UA exhibited the strongest positive effect (**Fig. 3f**, **Extended Data Fig. 5f**). Notably, the impact of upstream AUG (uAUG) on TE became increasingly negative as it approached the start codon, whereas CG showed a progressively positive effect. Albeit smaller in magnitude, most of the effects seemed to be maintained in the 3′ UTR, especially for regions proximal to the stop codon, suggestive of a position-dependent modulatory role for downstream AUGs and other dinucleotides. Taken together, these results establish that RiboNN captures the positional effects of nucleotide compositions along the entirety of the mRNA.

mRNAs with high TE are typically enriched for optimal codons^18,67^. To ascertain whether RiboNN has also learned this property, we reiterated our insertional analysis using 61 codons (excluding the 3 stop codons) inserted into the first reading frame along the length of each ORF. Similar to our previous findings, the model attributed substantially different effect sizes to codons depending on their position along the ORF, with the greatest predicted effects occurring near the start codon (**Fig. 3g**, **Extended Data Fig. 5g**). GCU (alanine), GGU (glycine), GAU (aspartic acid), and AAC (asparagine) exhibited the strongest positive effects on TE; conversely, AGG, AGA (arginine), UCA (serine), and UUA (leucine) showed the most negative impact^42^.

Based on the insertional analysis, we calculated the mean codon influence (*i.e.*, across the ORF) on TE for each of the 61 non-stop codons and observed a strong correlation between the scores derived from human and mouse RiboNN models (r=0.95, rho=0.95; **Fig. 3h**), indicating evolutionary conservation of predicted codon function on TE and the models’ ability to learn these reproducibly from completely independent datasets. Given the close link between codon usage and other aspects of RNA metabolism, we compared the correlation of RiboNN-based codon influence scores with several other metrics. We observed a strong positive correlation with mean codon stability coefficients^47^, which measure the association between codons and mRNA stability (**Fig. 3i**); a moderate negative correlation to propensity of ribosomes to have open A-sites^62^, which is indicative of ribosomes in the pre-accommodation state and hence slower elongation (**Fig. 3j**); and a moderate positive correlation with tRNA abundance^63^, which measures the availability of the cognate tRNA in the cellular pool (**Fig. 3k**). The correlations persisted when the scores of codons encoding the same amino acid were averaged, although no obvious trend existed with respect to hydropathy or charge of the amino acid (**Extended Data Fig. 6**). These findings underscore the complex interplay of multiple mechanisms that determine the fate of mRNAs in protein production.

### Predicting translational outcomes for therapeutically delivered mRNA sequences and genetic variants

Given RiboNN’s strong performance in predicting TE for endogenous mRNAs, we assessed its ability to generalize to orthogonal measures of TE and predict the impact of mRNA sequence variants on TE. Mean ribosome load, measured via polysome profiling, serves as an alternative metric of the translation rate of specific mRNAs, whether endogenous or therapeutic. Unlike ribosome profiling, mean ribosome load can differentiate translation differences between multiple RNA transcript isoforms of a given gene^21,68,69^. RiboNN, which was modeled on the full length of mRNAs, can be easily adapted to predict such isoform-specific TEs. The HEK293T RiboNN model demonstrated r^2^=0.34 and r^2^=0.69 between predicted TEs and mean ribosome loads measured for endogenous transcripts, which is within the realm of the reproducibility of measurement between labs (r=0.73; **Fig. 4a**). These results indicate that our model effectively captured the relationships between isoform diversity and translational regulation.

**Fig. 4.**
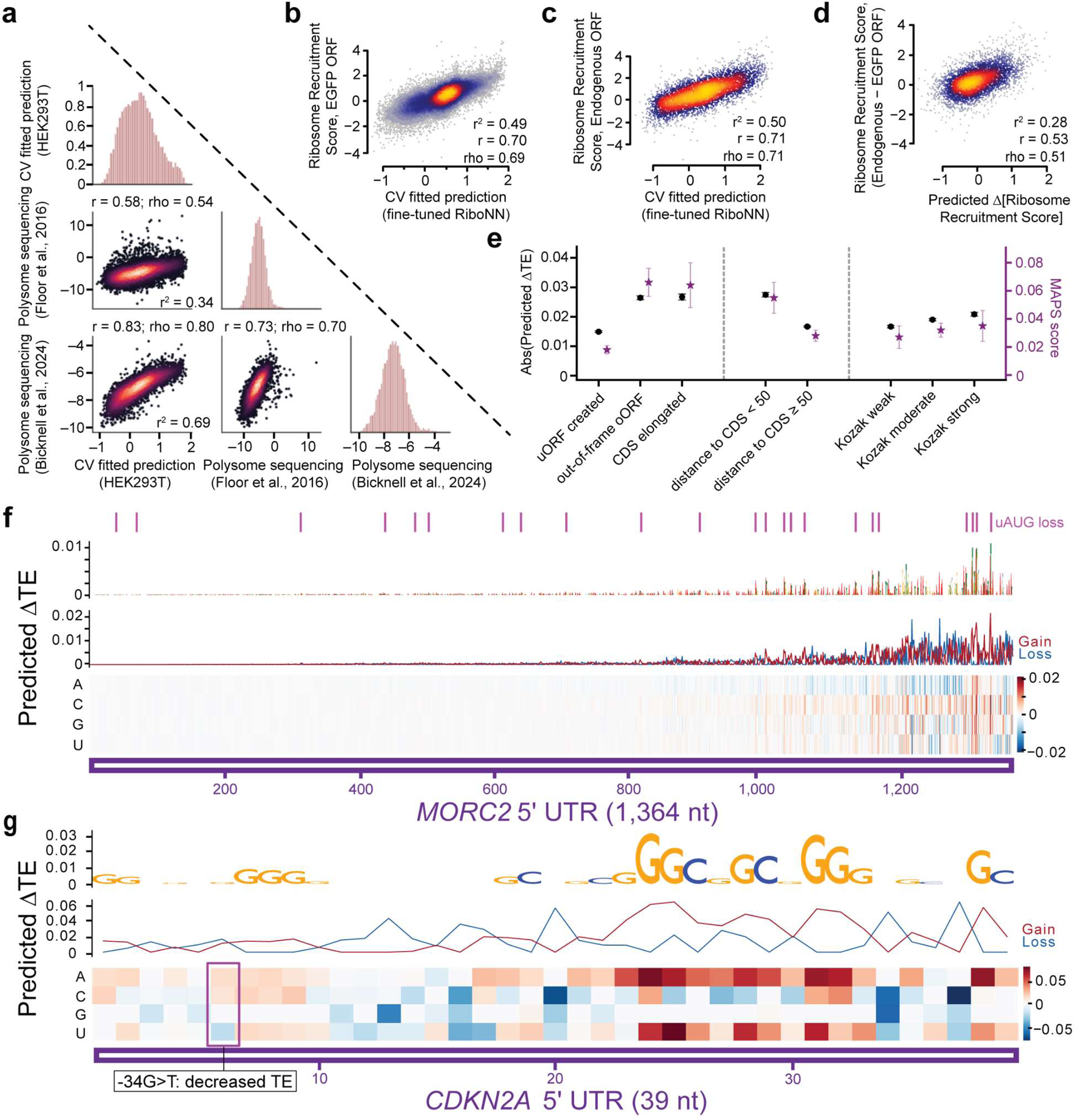
RiboNN predicts the impact of RNA modifications, genetic variants, and reporter constructs on translation. **a,** Comparison of HEK293T-predicted TEs relative to mean ribosome load as measured by polysome profiling^21,72^. **b-d,** Performance of RiboNN fine-tuned to predict the ribosomal recruitment score (*i.e.*, association of the 80S ribosomal subunit) to a panel of m^1^Ψ-modified 5′ UTRs linked to EGFP (**b**), their corresponding endogenous ORFs (**c**), or the paired difference between the endogenous and EGFP ORF (**d**)^61^. The Pearson (r) and Spearman (rho) correlation coefficients between each pair of measurements is also shown. **e,** Relationship between the observed strength of negative selection of uAUG-associated point mutations, as measured by the mutability adjusted proportion of singletons score^71^, and the RiboNN-predicted effect size. uAUG mutations were binned into categories based on the type of ORF created, distance to CDS start position, and association to Kozak consensus sequences of varying strength^71^. Error bars represent confidence intervals calculated using bootstrapping^71^. **f-g,** *In silico* mutagenesis results of two 5′ UTR regions of *MORC2* (**f**) and *CDKN2A* (**g**). “Gain” alludes to a predicted increase in TE for the mutation, while “Loss” refers to the opposite. Plotted are the sequence logo of the wild-type sequence, with nucleotide height proportional to the average predicted gain in TE (top row); the maximum predicted gain or loss across all possible mutations at each position (middle row); or a heatmap of all predicted TE changes for all possible mutations (bottom row). Positions of wild type uAUG are highlighted in purple at the top. The known disease associated variant is boxed.

In addition to endogenous mRNAs, polysome profiling has been used to measure translation from reporter constructs and base-modified mRNAs, as these can significantly influence protein output^70^. We next tested RiboNN’s ability to predict mean ribosome load in a massively parallel reporter assay dataset which assessed the function of thousands of 5′ UTRs in parallel^14^. Although RiboNN was never trained on polysome profiling or reporter data, its predicted TEs were still associated with mean ribosome load, achieving an r^2^ of 0.14−0.15 for reporter mRNAs without modified bases and an r^2^ of 0.09−0.10 for reporter mRNAs with either Ψ-modifi-ed or N1-methylpseudouridine (m^1^Ψ)-modified nucleotides (**Supplementary Fig. 9**). It’s poorer performance relative to Optimus^14^ and FramePool^16^ was not surprising given that the latter two were trained directly on these datasets. Thus, to more fairly benchmark RiboNN against these models, we benchmarked all three on a third-party dataset which measured ribosome recruitment scores for mRNAs with m^1^Ψ-modified 5′ UTRs linked to different ORFs^61^. RiboNN was weakly predictive of this data (r^2^=0.17 for 5′ UTRs linked to EGFP; r^2^=0.19 for 5′ UTRs linked to endogenous ORFs; **Supplementary Fig. 10a-b**). Leveraging the paired measurement of endogenous ORF and EGFP, we observed r^2^=0.11 between changes in TE and changes in ribosome recruitment scores resulting from swapping the ORFs (**Supplementary Fig. 10c**). In contrast, prior 5′ UTR-based models could not predict differences among 5′ UTRs linked to different ORFs. Depending on the ORFs tested, Optimus^14^ performed 2−7-fold worse (r^2^=0.08 on EGFP ORF; r^2^=0.03 on endogenous ORFs) than RiboNN; likewise, FramePool^16^ performed 2−5-fold worse (r^2^=0.08 on EGFP ORF; r^2^=0.04 on endogenous ORFs) (**Supplementary Fig. 10d-g**). Given the limited predictive power of all three models, we examined whether fine-tuning our RiboNN model on these data could improve performance (*i.e.*, via transfer learning). Indeed, fine-tuning it improved performance by 2−3-fold relative to the original model (r^2^=0.49 for 5′ UTRs linked to EGFP; r^2^=0.50 for 5′ UTRs linked to endogenous ORFs; r^2^=0.28 for predicting ORF-dependent 5′ UTR effects; **Fig. 4b-d**). These improvements could not be explained by merely re-training RiboNN from scratch, illustrating the power of transfer learning in this context (**Supplementary Fig. 10h**). Collectively, these findings underscored RiboNN’s ability to integrate information from both 5′ UTR and ORF regions while predicting the translational regulation of therapeutically relevant mRNA.

Utilizing the entire mRNA sequence enables the examination of how differences in sequence, including disease-associated variants, influence TE at single-nucleotide resolution. Given that 5′-UTR variants that generate or disrupt uORFs can lead to disease and are key *cis*-regulators of tissue-specific translation^71^, we first assessed RiboNN’s ability to predict the impact of uAUG-associated point mutations. The RiboNN-predicted effect size had a strong association with the strength of negative selection, as indicated by the mutability-adjusted proportion of singletons score^71^ (**Fig. 4e**). Variants creating uAUGs that result in overlapping open reading frames (oORFs) or elongated CDSs exhibited a significantly higher impact on the TE of downstream protein-coding genes; moreover, uAUGs generated within 50 nt of the CDS had a greater effect size than those created further upstream (**Fig. 4e**). The effect size is slightly elevated if uAUG-creating variants arise in the context of strong Kozak consensus sequences relative to moderate or weak ones (**Fig. 4e**). These findings reveal that RiboNN learned positional and contextual features of uAUGs, both in function and evolutionary constraint.

Next, we conducted *in silico* mutagenesis on the 5′ UTR regions of several disease-associated genes. *MORC2*, a gene implicated in Charcot-Marie-Tooth disease^73^, has a long 5′ UTR region with a large number of uAUGs. Reinforcing earlier results (**Fig. 4e**), RiboNN predicted that loss-of-function mutations in CDS-proximal uAUGs would have a greater effect size relative to distal uAUGs (**Fig. 4f**). For the gene *RDH12*, associated with inherited retinal disease, RiboNN successfully predicted the negative impact of a uAUG-creating SNP (−123C>T), which had been experimentally validated to reduce translation^74^ (**Extended Data Fig. 7a**). Additionally, the gene *CDKN2A* has a reported G>T mutation at base −34 in its 5′ UTR that creates a uAUG reported to decrease translation, leading to predisposition to melanoma^75^. RiboNN consistently predicted decreased TE for this variant (**Fig. 4g**). The ability of RiboNN to correctly predict the impact of TE of variants extended beyond those associated with uAUGs. For example, the SNPs −127C>T and −9G>A in the 5′ UTR of the *ENG* gene, associated with hereditary hemorrhagic telangiectasia, have been reported to reduce the expression levels of *ENG*^76^, consistent with the decreased TE predicted by RiboNN (**Extended Data Fig. 7b**). For *FGF13*, a gene associated with congenital intellectual disability, the −32C>G mutation reduces translation^77^. RiboNN also predicted a negative effect of this SNP on TE, and indicated that a C>A mutation at the same position might have an even greater impact on TE (**Extended Data Fig. 7c**). However, for SNP −94G>A in *BCL2L13*, RiboNN predicted an increase in TE, contrary to the reported decrease in protein expression^78^ (**Extended Data Fig. 7d**). These results suggest that RiboNN could offer an additional form of evidence to infer the regulatory impact of SNPs on disease-associated genes.

### RiboNN learns a common language governing mRNA stability, translational regulation, and localization

Given the strong positive correlation between the RiboNN’s mean codon influence on TE and the previously estimated codon influence on mRNA stability (**Fig. 3i**), we further assessed the relationship between TE and mRNA stability. Indeed, both the predicted and experimentally measured mean TE as well as mRNA stability from a previous study^7^ were positively correlated in humans and mice (r>0.31, rho>0.32; **Fig. 5a, Supplementary Fig. 11**). Similar patterns were also observed between mRNA stability, polysome profiling, and PTR data, with the exception of the PTR dataset^30^ previously observed to be an outlier (**Supplementary Fig. 11a**, **Fig. 1c**). Consistent with the predicted underlying role of codons influencing both TE and stability, mean TE (as predicted by RiboNN) was positively correlated with mRNA stability (r=0.38, rho=0.36; **Fig. 5b**); conversely, mRNA stability (as predicted by Saluki^7^ was positively correlated with TE (r=0.40, rho=0.40; **Fig. 5c**). Taken together, these results suggest an interconnectedness between mRNA stability and translational regulation that can be learned by sequence-based machine learning models from diverse and independent datasets.

**Fig. 5.**
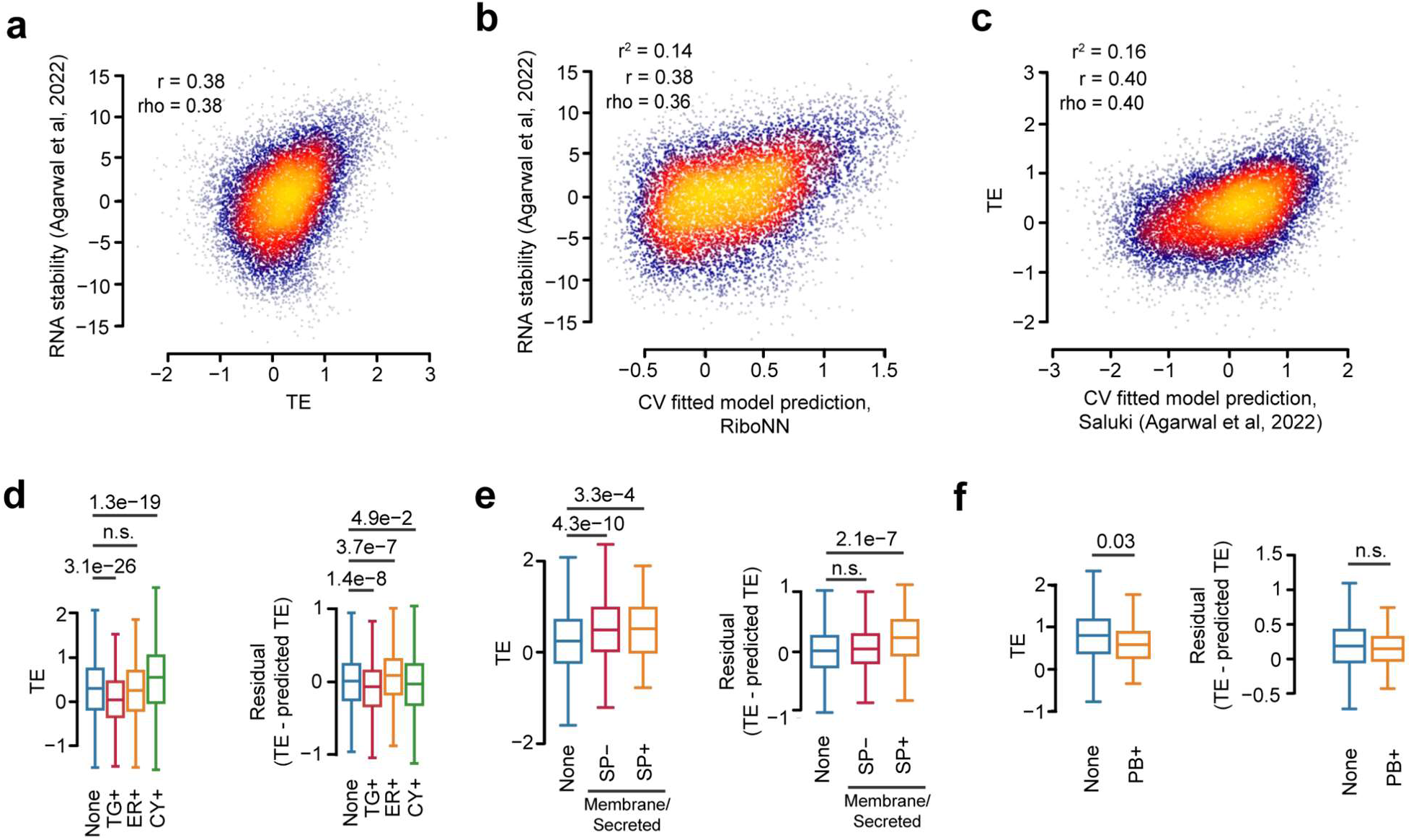
Interrelationships between mRNA translation, turnover, and subcellular localization. **a-c,** Scatter plots showing the relationship between mean TE and mRNA stability^7^ (**a**), predicted mean TE and mRNA stability (**b**), and predicted stability and mean TE (**c**). Pearson (r) and Spearman (rho) correlation coefficients are also indicated. **d-f,** Boxplots of TE (left panel) and residual TE (*i.e.*, representing the difference between TE and the predicted TE, right panel) for mRNAs binned according to their subcellular localization. Shown are the distributions for mRNAs encoding non-membrane (excluding secreted) proteins that are enriched in TIS granules (TG+), rough endoplasmic reticulum (ER+), or cytosol (CY+)^79^ (**d**); mRNAs encoding membrane or secreted proteins, with or without predicted signal peptides (SP+/–)^80^ (**e**); or mRNAs enriched in cytosolic processing bodies (P-bodies)^81^ (**f**). p-values were computed by comparing the behavior of mRNAs localized to the specified compartment relative to those not localized (*i.e.*, labeled “None”) using a two-sided Mann-Whitney test adjusted with a Bonferroni correction.

mRNAs localized to certain subcellular compartments, such as the endoplasmic reticulum (ER) membrane, tend to be differentially translated^79,82^. We sought to evaluate these findings in the context of our predictive model, assessing both TEs and their associated residuals (mean TE – predicted mean TE) for mRNAs localizing to different compartments. For mRNAs encoding non-membrane (excluding secreted) proteins, we observed a significantly higher residual TE for ER-enriched mRNAs; additionally, cytosolically enriched mRNAs exhibited a higher TE, although this signal was largely explained by the model (**Fig. 5d**). When considering mRNAs encoding both non-membrane and membrane or secretory proteins, a higher TE was observed for ER-enriched mRNAs (p<0.01, data not shown). This is consistent with the result that mRNAs encoding membrane or secreted proteins tended to have higher TE, even for those lacking a signal peptide sequence (**Fig. 5e**). Nevertheless, membrane/secreted proteins harboring a signal peptide possessed a strongly positive residual on average (**Fig. 5e**), indicating that RiboNN was unable to model the association between signal peptides and TE. This was unsurprising as the model was trained on ∼10K mRNA sequences and the number of mRNA sequences encoding signal peptides is combinatorially explosive.

Given past work finding a relationship between mRNA stability and localization^18^, we evaluated whether unexplained variation in TE from RiboNN’s predictions could also be linked to mRNA localization. Since less stable mRNAs tend to be translationally repressed and enriched in mRNA processing bodies^81^ (P-bodies), we expected that mRNAs enriched in P-bodies to have lower mean TE compared to other mRNAs. This indeed appeared to be the case (**Fig. 5f**); however, there was no difference in the residual between mRNAs enriched in P-bodies (“PB+”) and others (“None”), indicating that the model already learned that mRNAs enriched for localization to P-bodies was associated with differential TE (**Fig. 5f**). Collectively, our results thereby establish a common language governing mRNA decay, translational regulation, and subcellular localization.

## DISCUSSION

In this study, we developed deep learning models that utilize entire mRNA sequences to predict TE. These models were trained using data synthesized from thousands of ribosome profiling and matched RNA-seq experiments across >140 human and mouse cell types. Our models explain over 70% of the variation in TE in specific cell lines, achieving a mean r^2^ across cell types of 0.62. This represents a 1.3 to 4.4-fold performance improvement relative to previously developed models in mammals, which achieved a maximum r^2^ of 0.46 (range from 0.14 to 0.46)^16,24,25,83^. Furthermore, unlike earlier efforts which were limited to a few cell types, our approach enabled the development of models for a substantially larger and more diverse set of cell types.

Recent research has primarily relied on reporter constructs to dissect regulatory elements of translation^14,15,59,84^. Due largely to technological limitations, such experiments employ easily detectable and fixed coding regions, such as GFP, attached to variably engineered 5′ UTRs, and are typically limited to one or few cell types. Critically, these reporter constructs lack the full complement of proteins that normally accompany endogenous mRNAs throughout their lifecycle^85^, which influences RNA metabolism^86^. Consequently, predictive models based on reporter assays offer limited insights into the translation of endogenous mRNAs, explaining less than 25% of variation in their TE^16,25^. In contrast, our model demonstrates superior performance in predicting the translation of endogenous mRNAs and also appears to predict the behavior of therapeutic RNAs^61^.

Our predictive modeling approaches are particularly valuable as they provide a quantitative assessment of factors determining TE. By analyzing the position and identity of sequence elements, we were able to ascertain their relative importance in making accurate predictions. Our model highlights the dominant influence of 5′ UTRs and coding sequences in determining TE. The nucleotide compositions of 5′ UTRs heavily influenced the prediction of TE. Short, AU-rich sequences were generally associated with higher TE, whereas the impact of GC-rich sequences was negative but position-dependent. Intriguingly, recent massively parallel reporter assays conducted in both zebrafish and human cells, utilizing different readouts to measure translation, have identified a similar pattern^59,61^. This concordance suggests that these particular regulatory features observed in reporter constructs are reflective of those in endogenous transcripts.

RiboNN also learned the well-established role of uAUGs in repressing the translation of the main coding sequence^14,60,78,87^. Specifically, a shorter distance between the uAUG and the start codon was associated with a reduced TE of the main coding sequence, consistent with the depletion of uAUGs near CDS start sites^83^. Furthermore, uAUGs closer to the start codon are more likely to produce overlapping ORFs. Such overlapping ORFs, which are under more stringent selective pressure in human populations^71^, tend to inhibit the TE of the main CDS more than uORFs entirely contained within the 5′ UTR, which may allow for reinitiation following uORF translation termination^60^.

In addition to learning the well-established role of uAUGs, our model unexpectedly predicts that downstream AUGs in 3′ UTRs reduce TE, particularly when close to the stop codon. Readthrough of stop codons can lead to C-terminal extensions, which decrease protein abundance^88^. The underlying mechanisms likely involve both proteasomal degradation^88,89^ and reduced translation due to ribosome stalling^90,91^. Alternatively, downstream AUGs can be translated due to inefficient recycling of terminating ribosomes that subsequently reinitiate^92^. Although the impact of such events on the TE of the main ORF remains incompletely understood, a recent study suggested that translation of downstream ORFs can act as translational activators^93^. While our findings might appear to contradict this finding, it is conceivable that there is a distance-dependent relationship, where AUGs near stop codons are inhibitory due to their effects on recycling efficiency or readthrough, whereas ORFs positioned further downstream could have activating effects. Although our models detect specific signals in 3′ UTRs, particularly near the stop codon, overall, RiboNN predicts that 3′ UTRs generally have a minimal impact on TE. Our results do not imply that 3′ UTR-dependent regulation is unimportant for specific genes^94^ or particular contexts such as in early vertebrate development^95,96^. However, the overall contribution of 3′ UTRs to translation control is likely limited, consistent with several transcriptome-wide analyses ^31,97^.

A major finding from our study is the dominant influence of the coding sequence on TE predictions. Particularly, sequences proximal to the N-termini were found to be about twice as important in determining TE, a feature learned by RiboNN independently from both mouse and human datasets. Interestingly, recent work using reporter constructs and single-molecule analyses suggested that the identity of amino acids in early coding regions can affect protein synthesis efficiency, potentially through mechanisms related to translation elongation^44^. While the N-terminus-proximal codons were more important at a per-residue level, the identity of codons across the entire CDS contributed to TE predictions. Factors such as the charge of the nascent polypeptide in the exit tunnel of the ribosome^98,99^, the pairs of codons in the decoding center^42,100^, and availability of charged tRNAs corresponding to specific codons^101^ have all been linked to altered translation elongation. Despite these mechanisms that can alter decoding rates, there is debate over whether the average elongation rate across different mRNAs varies significantly^102,103^. Critically, recent studies implicate codon usage in modulating initiation efficiency through differences in ribosome decoding rates^104,105^. Given the importance of the entire CDS for the accuracy of RiboNN, our results suggest that both codon and amino acid compositions are critical for determining the TE of endogenous mRNAs.

Translation elongation dynamics have emerged as an important contributor to mRNA stability as well^17,18,45–49^. The codon-specific effects identified by RiboNN in predicting TE closely mirror their impact on mRNA stability. For instance, the codons AGA and AGG, which were found to exert significant mRNA-destabilizing effects^7,106^, also negatively impact TE, as inferred by RiboNN. Additionally, during the maternal-to-zygotic transition, mRNAs enriched with codons that enhance mRNA stability also show higher TE^17^. However, the relationship between translation and mRNA decay remains debated^107^, as increased TE and ribosome flux can also facilitate mRNA decay, which would predict a negative correlation between the two^72^. Specifically, slower elongation rates may result in mRNA degradation through either transiently slowed ribosomes^108,109^ or ribosome collisions, which can activate the ribosome quality control pathway^110^. While these mechanisms have been primarily explored using reporter constructs, recent studies have also demonstrated its relevance to endogenous transcripts^111^. Detailed investigation into the translation-dependent and independent contributions to mRNA decay remains an active area of research^112^. Future studies are likely to uncover condition-specific effects on mRNA stability that vary with TE.

A potential limitation of our work is that it solely considers the primary sequence to predict TE. In our analyses using the LGBM and RiboNN models, the inclusion of several secondary structure-related features did not enhance performance. This might be explained by several possibilities: i) the primary sequence itself is highly predictive of secondary structure, potentially capturing these influences implicitly, ii) prior results may have overstated the importance of RNA structure because they did not appropriately account for nucleotide composition^43^, or iii) the list of structural motifs and features we computed, based on predicted free energy, do not accurately reflect the true secondary structures of these RNAs. Considering this last point, developing more precise secondary structure features could lead to further improvements in prediction accuracy. However, an independent analysis focused on RG4s, considered to be the most stable RNA structure that could block ribosome scanning^113^, suggested a weak but detectable translational inhibition for putative RG4-containing transcripts relative to RiboNN’s predictions. These findings would require further validation considering the counteracting evidence that RG4s are globally unfolded in mammalian cells^114^.

Another avenue for improvement could involve providing RiboNN with explicit knowledge of protein sequences. Including amino acid composition information improved the performance of the LGBM model, and our analyses revealed systematic bias in predicted TE for proteins harboring signal peptides. Thus, a deep learning model that accesses both nucleotide and amino acid sequence (*i.e.*, or summarized protein-based information), may further enhance TE prediction. Nevertheless, since our models currently explain 62% of the variability in mean TE across a wide array of cell types, we can establish an upper bound on the impact of such features. This estimate is likely conservative, as some portion of the unexplained variance in these measurements is attributable to measurement error.

We would also like to note that TE, as defined in our study and typically used in the literature, does not equate to the rate of protein synthesis; rather, it reflects differences in ribosome occupancy relative to mRNA abundance. While recent work with reporter constructs suggested that increased ribosome load may not linearly relate to protein output^72^, both our work and previous studies^32,115^ indicate that TE is positively associated with protein abundance and synthesis rates for endogenous transcripts. Theoretical models of translation also support the general positive relationship between protein synthesis and TE^56,116^.

Overall, RiboNN achieves state-of-the-art prediction of TE in humans and mice, elucidating key principles that underpin accurate predictions, including the relative importance of various molecular aspects. These predictive models distill our knowledge into a coherent framework and have the potential to advance bioengineering applications. Significantly, RiboNN has the ability to generate functional predictions on genetic variants in the human population, giving insight into the mechanisms constraining molecular evolution and underpinning genetic diseases. Overall, these advancements have far-reaching implications for both genetic diagnostics as well as the design and optimization of mRNA and gene therapies, positioning our model at the forefront of these rapidly evolving domains. Looking ahead, we anticipate that future work will employ multi-modal approaches to simultaneously predict all facets of gene expression—RNA abundance, stability, and translation—from primary mRNA sequence, given the interconnectedness of these phenomena.

## ACKNOWLEDGMENTS

We thank Ian Hoskins (UT Austin) for the code and data to generate secondary structure features, and Milad Miladi (Sanofi) for providing critical feedback. We thank Carson Thoreen and Wendy Gilbert (Yale University) for sharing their data prior to publication. Research reported in this publication was supported in part by the National Institute Of General Medical Sciences of the National Institutes of Health under Award Number R35GM150667 (C.C.). This work was also supported by the National Institutes of Health grant [HD110096], and the Welch Foundation grant [F-2027-20230405] (C.C.). C.C. was a CPRIT Scholar in Cancer Research supported by CPRIT Grant [RR180042].

## AUTHOR CONTRIBUTIONS

D.Z. trained RiboNN models, validated model predictions with public datasets, and contributed to model interpretation. J.W. interpreted RiboNN, performed comparisons between TE and third-party measurements, and analyzed genetic variant data. L.P. trained and interpreted classic ML models. Y.L. helped synthesize the data compendia and developed the compositional approach to calculate TE. F.M., C.C., and V.A. supervised the study. C.C. and V.A. conceptualized and designed the study.

## CODE AND DATA AVAILABILITY

Code, pre-trained models, and data are planned for public release upon successful review of this article. Our classic ML model code is available at: https://github.com/CenikLab/TE_classic_ML.

## SUPPLEMENTARY TABLES

**Supplementary Table 1. Feature sizes, sequences, CV folds, and TEs of human and mouse genes.** The principal splice isoforms for human and mouse genes were downloaded from APPRIS v2^117^. The CV folds reported were used to split training and test sets. The TEs of transcripts with an average coverage <0.1x were set to NA. The mean TEs were calculated across the cell types for each transcript while ignoring NA values.

**Supplementary Table 2. Feature sizes, sequences, CV folds, and TEs predicted by the human and mouse RiboNN models.** The principal isoforms for human and mouse genes were downloaded from APPRIS v2^117^. Predicted results are reported for the multitask and single-task RiboNN models (described in **Fig. 3d**). For transcript/cell type combinations in which the TE is NA in the training data, the predicted TEs were set to NA.

## METHODS

### Generation of human and mouse TE compendia

To calculate cell-type-specific TEs, we initially selected 1,282 human and 995 mouse ribosome profiling datasets with matched RNA-seq data. These were screened for a series of quality control steps to retain high-quality samples. Quality control criteria included ensuring average transcript coverage exceeded 0.1X and reads mapping to CDS constituted more than 70% of the total. The remaining 1,076 human and 835 mouse ribosome profiling samples were further processed using the winsorization method to minimize the impact of PCR bias^52^. Genes with sufficient counts per million (CPM > 1 in more than 70% samples) of RPFs were retained, and transcripts without poly(A) tails were removed. Experimental variables, such as the inclusion of elongation inhibitors, can lead to technical artifacts, manifesting as increased RPF density around start and stop codons^118^. To mitigate such biases, we only considered RPFs whose 5′ end mapped either after the first 10 nts or before the last 35 nts of the CDS. These RPFs were summed to determine the CDS count for each transcript^51^. An identical counting method was used for RNA-seq data. Total CDS counts for both RNA-seq and ribosome profiling were normalized using a centered log-ratio. TE was defined as the residual obtained from a compositional linear regression, for each transcript in each sample^52^. For each transcript, if either the RNA-seq or ribosome profiling read count was 0 in all samples from a specific cell line, we assigned NA to its TE in the corresponding cell line. Finally, we calculated the average TE for each transcript in each cell line across all samples.

### Features considered in classical machine learning models

The length features included the log_10_ of the 5′ UTR, CDS, 3′ UTR, and total transcript lengths. Nucleotide frequency included the percent composition of the 5′ UTR, CDS, 3′ UTR and full sequence. Codon and amino acid frequencies were calculated as the percentage within the CDS, and included annotated stop codons. K-mer frequencies (for k-mers of size two through six) were computed separately for each region and normalized by the total k-mer count. Additional feature classes included the frequency of each nucleotide in the wobble position of all codons, a one-hot encoding of the nucleotide identity surrounding the start codon (at the −3, −2, −1, +4, and +5 positions), the counts of 20 dicodons found to affect TE in yeast^42^, and several secondary-structure-related metrics. To capture secondary structure, sequences for the 5′-most 60 nt of the transcript and a 60 nt window centered on the start codon (*i.e*., last 30 nt of the 5′ UTR and first 30 nt of the CDS) were extracted from the APPRIS v2 primary transcript references^117^. If the 5′-UTR length was <30 nt, the first 60 nt of the transcript were used instead. Secondary structure features were enumerated in these regions using seqfold v0.7.17 (https://github.com/Lattice-Automation/seqfold, https://zenodo.org/records/7986470) at a temperature of 37°C. These features were the min ΔG, number of hairpins, number of loops, number of bifurcations, number of bulges, max stem length, max loop length, and position of the first stem. Hairpins with a stem length <3 or loop length >10 were not enumerated. Some of the sequence-derived and biochemical features used previously^7^ were also tested separately and in combination with the above sequence features. The sequence-derived features include G/C content, intron length, ORF exon junction density, predictions for mammalian microRNA targets, and the average binding score of mammalian RNA-binding proteins (RBPs). The biochemical features include the measured mRNA half-life; number of CLIP, eCLIP, and PAR-CLIP peaks of various RBPs; and the enrichment of RBP binding relative to a control IP measured by RIP-seq^7^.

### Classical machine learning model benchmarking

The lasso, elastic net, random forest (scikit-learn v1.0.2)^119^, and LGBM (lightgbm v3.2.1)^120^ regression models were trained using 10-fold CV. Performance was measured as the mean of the r^2^ values across held-out test folds. Throughout this study, r^2^ (i.e., the coefficient of determination) was computed as the square of the Pearson correlation. For lasso and elastic net, the training data was further split into 5 CV folds to find the optimal α (lasso and elastic net) and L1 ratio (elastic net) hyperparameters. The default hyperparameters given were used for LGBM, with the exception of the “gain” option for use with importance calculations. Random forest used the same number of trees and maximum leaf nodes as LGBM. Comparisons between model types (**Extended Data Fig. 3**) and feature sets (**Fig. 2a**) were deemed significant with one-sided, paired t-tests, adjusted by a Bonferroni correction. We measured feature importance as the sum total information gain across all LGBM tree splits using that feature, averaged across all folds. In **Fig 2b-c**, the importance was further averaged over all cell lines. To determine if a feature had a positive or negative effect on prediction, the Spearman correlation between the feature and cell-type-specific TE was used.

### RiboNN model architecture, training, and interpretation

The input mRNA sequences were aligned at the start codons, with the maximum 5′ UTR size set to 1,381 nt and the maximum combined CDS and 3′ UTR size to 11,937 nt. Sequences were padded at the 5′ and 3′ ends with “N”, and one-hot encoded (with ‘N’ encoded by a vector of four 0s). We added a fifth channel labeling the first nucleotide of each codon in the CDS^7^.

The architecture of RiboNN consisted of a Conv1D input layer, a “tower” of ten convolution blocks, and a head of 2-linear layers (**Extended Data Fig. 2**), with each convolution block including the following operations: i) layer normalization sandwiched by transpose actions, ii) ReLU activation, iii) 1D convolution with kernel width 5, iv) dropout, and v) max pooling with width 2. Overall, the model consisted of 250,382 learnable parameters. The output layer had one or multiple neurons for single-task and multitask learning, respectively.

Following Saluki’s training procedure^7^, we trained the RiboNN multitask model with the MSE loss function using the AdamW optimizer on batches of 64 randomly selected examples, a gradually decreasing learning rate between 0.001 and 0.0000001, beta1 of 0.9, and beta2 of 0.998. We clipped gradients to a global norm of 0.5. We used a dropout probability of 0.3 throughout. After each epoch, the model was evaluated on the held-out validation set in batches of 128 sequences. We trained each model for 200 epochs, saving checkpoints along the way. After 200 epochs, the model parameters from the checkpoint with the highest validation r^2^ were saved as the final model parameters. We trained the mouse and human models independently using a nested CV strategy. Specifically, we randomly split the full set of human or mouse transcripts into 10 folds of similar sizes and trained 9 models for each of the 10 held-out CV folds (using 9-fold CV on the inner folds), producing a total of 90 trained models. For each of the 9 models from the inner folds, we retained the top 5 models ranked based upon their validation r^2^ performance. When running RiboNN in “prediction” mode, we computed the mean of these 50 models to represent the ensemble prediction.

Transfer learning was implemented by replacing the linear head of our pre-trained multitask model with a new single-task 2-layer linear head. We froze all preceding layers and trained the new linear head for 50 epochs, followed by unfreezing all of the layers and training the entire network for another 150 epochs. To prevent circularity during transfer learning (e.g., use of the same gene in the tissue-specific TE test set as was considered in the multitask model training set), we fine-tuned the pre-trained multitask model on the matched train/test splits.

We used the saliency method^65^ within the PyTorch Captum library (version 0.6.0)^121^ to compute the attribution scores for each nucleotide of the input sequence with respect to the predicted mean TE. For each of the test sets from our 10-fold CV procedure, we averaged the attribution scores from the top 5 trained models.

To generate the metagene plot of attribution scores, we followed the methods established in prior work^7^.

### Insertional motif analysis with RiboNN

Using attribution scores as input, we ran TF-MoDISco-lite^66^ on each functional region (5′ UTR, ORF, and 3′ UTR) independently to identify the motifs most strongly influencing the predicted mean TE. Gradient correction was applied by subtracting the mean attribution score across four encoding channels^65^. The motifs were ranked based on the number of sequences (*i.e.*, seqlets) supporting the enrichment of each motif.

As performed in earlier work^7^, the insertional analysis was performed by dividing each functional region of a valid mRNA into 100 evenly spaced positional bins. Each k-mer examined (*i.e.*, the 16 dinucleotides and AUG) was inserted into one of these bins, replacing the reference sequence to maintain the mRNA’s original length. A valid mRNA was defined as one with a 5′ UTR length ≥100 nt, an CDS length ≥500 nt, and a 3′ UTR length ≥500 nt^7^. For each insertion, the predicted change in mean TE relative to the corresponding wild-type mRNA was recorded. To quantify the impact of each motif across diverse sequence contexts, the predicted changes in mean TE across all valid mRNAs were averaged for each of the 300 positional bins. Identical insertional analysis was performed for the 61 non-stop codons, except that each codon was inserted into the first reading frame of the ORF.

### Impact of uAUG-creating variants with RiboNN

As described in an earlier study^71^, we retrieved the list of variants that create uAUGs and selected the canonical transcript based on the gnomAD v2 annotation^122^ for each gene for further analysis. For each uAUG-creating variant considered, we verified that its gene name matched the list of canonical transcripts and that the distance from each uAUG variant to the start of its CDS was accurately annotated. This led to a set of 15,184 uAUG variants which were categorized into two groups based on their effects and contexts as previously annotated^71^. The effect group was comprised of variants that create out-of-frame oORFs (n=2,784), elongate the CDSs (n=1,350), or generate uORFs (n=9,263). The context group included variants located at a distance of ≥50 nt from the CDS (n=11,113), <50 nt from the CDS (n=2,284), or associated with a strong (n=2,237), moderate (n=6,559), or weak (n=4,601) Kozak consensus sequence. To assess the impact of each variant on TE, we recorded the change in predicted TE relative to the wild-type mRNA reference sequence. The confidence intervals were calculated using bootstrapping as described^71^.

### *In silico* mutagenesis analysis of disease genes with RiboNN

We performed *in silico* mutagenesis analysis^7^ on the 5′ UTR regions of genes associated with various diseases to predict the impact of genetic variants on TE. For each nucleotide position, we substituted the reference nucleotide with each of the three possible alternative alleles, and computed the predicted ΔTE.

### Subcellular localization analysis

Based on prior results^79^, we categorized 5,884 non-membrane (excluding secreted) protein-encoding mRNAs as enriched in TIS granules (TG+, n=1,086), the rough ER (ER+, n=745), the cytosol (CY+, n=1,299), or exhibiting no apparent localization (2,754). For our analysis of P-body-enriched mRNAs, we examined a total of 1,636 mRNAs^81^, of which 93 exhibited P-body enrichment based on prior results^81^. P-values from Mann-Whitney-Wilcoxon test two-sided with Bonferroni correction were performed to show statistical significance.

