## Supplemental Figures for "Predicting the translation efficiency of messenger RNA in mammalian cells"

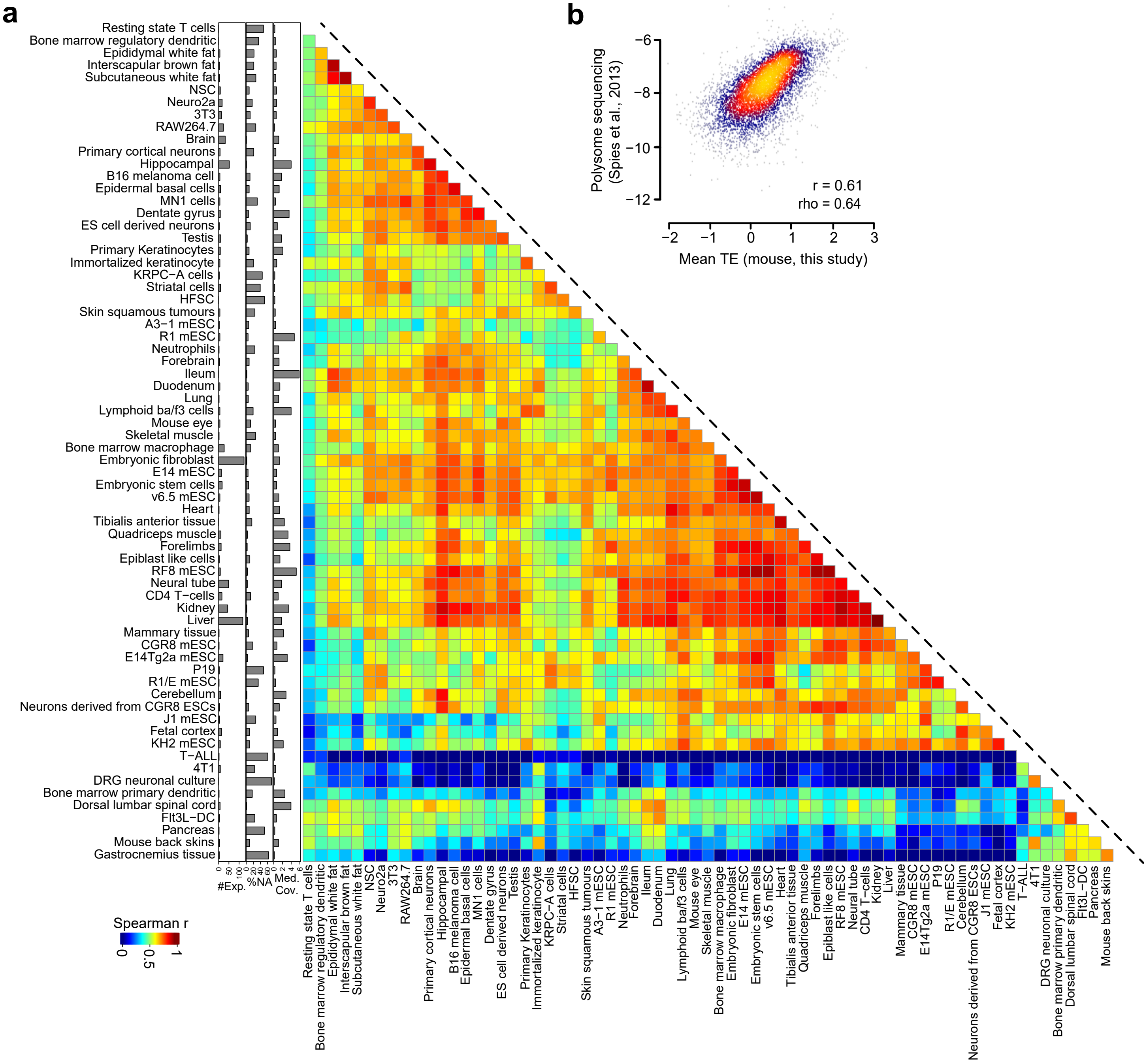

**Extended Data Fig. 1. Intercomparison of mouse cell types. a,** This panel is the same as that of **Fig. 1b** except displays results for 68 mouse cell types. **b,** Scatter plot to show the correlation between mouse mean TE from this study and ribosomal loading measured in 3T3 cells[1]. Pearson (r) and Spearman (rho) correlation coefficients are indicated.

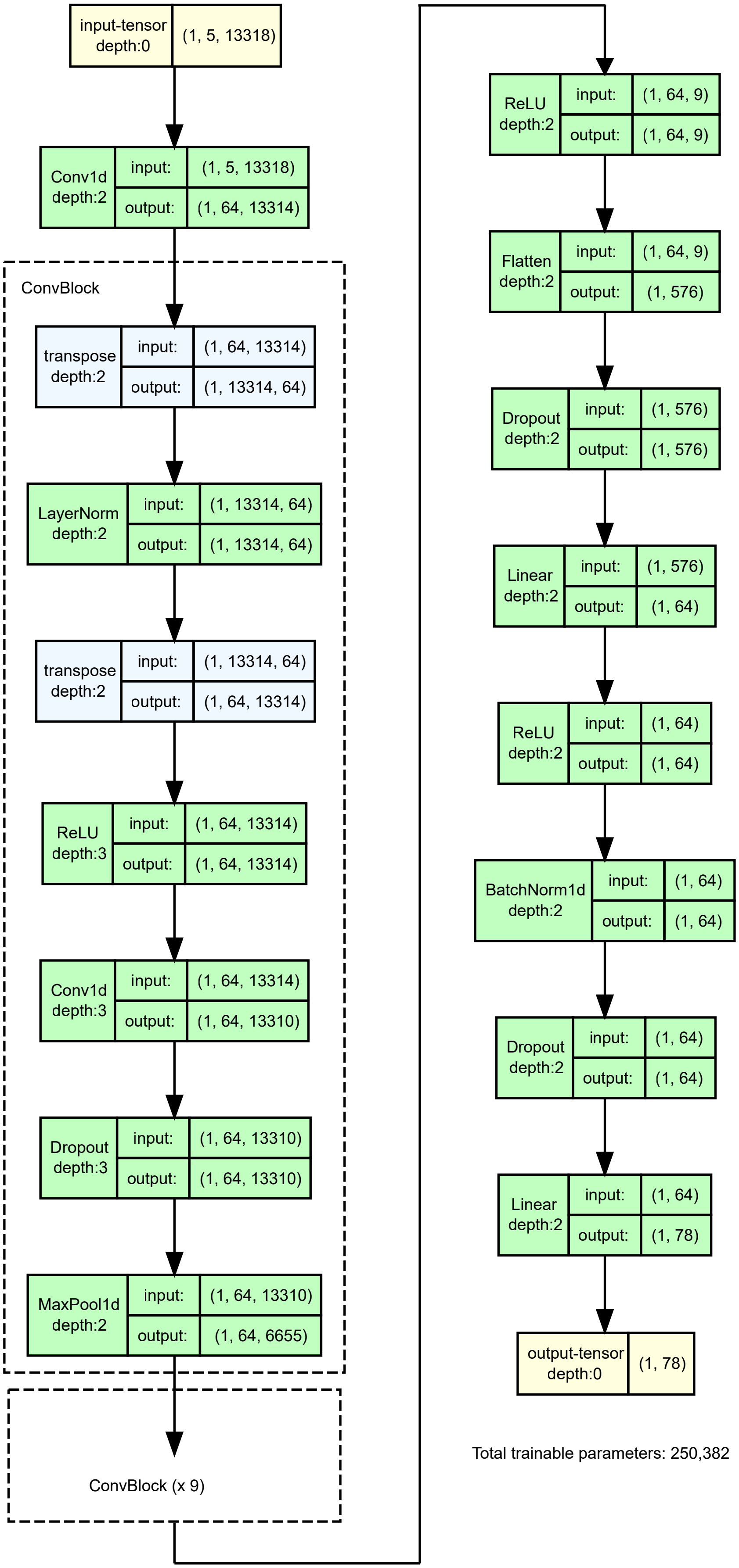

input-tensor depth:0 (1, 5, 13318)
Conv1d depth:2 input: (1, 5, 13318) output: (1, 64, 13314)
ConvBlock
transpose depth:2 input: (1, 64, 13314) output: (1, 13314, 64)
LayerNorm depth:2 input: (1, 13314, 64) output: (1, 13314, 64)
transpose depth:2 input: (1, 13314, 64) output: (1, 64, 13314)
ReLU depth:3 input: (1, 64, 13314) output: (1, 64, 13314)
Conv1d depth:3 input: (1, 64, 13314) output: (1, 64, 13310)
Dropout depth:3 input: (1, 64, 13310) output: (1, 64, 13310)
MaxPool1d depth:2 input: (1, 64, 13310) output: (1, 64, 6655)
ConvBlock (x 9)
ReLU depth:2 input: (1, 64, 9) output: (1, 64, 9)
Flatten depth:2 input: (1, 64, 9) output: (1, 576)
Dropout depth:2 input: (1, 576) output: (1, 576)
Linear depth:2 input: (1, 576) output: (1, 64)
ReLU depth:2 input: (1, 64) output: (1, 64)
BatchNorm1d depth:2 input: (1, 64) output: (1, 64)
Dropout depth:2 input: (1, 64) output: (1, 64)
Linear depth:2 input: (1, 64) output: (1, 78)
output-tensor depth:0 (1, 78)
Total trainable parameters: 250,382

**Extended Data Fig. 2. Visualization of the RiboNN model architecture.** Shown is a layer by layer graph of the RiboNN architecture, with input/output dimensions labeled for each layer. The ConvBlock in the broken-line box was applied 10 times in total to compress the sequence length. Light yellow nodes reflect input/output tensors, light blue nodes reflect functions, light green nodes reflect modules, and numbers in parentheses reflect tensor dimensions.

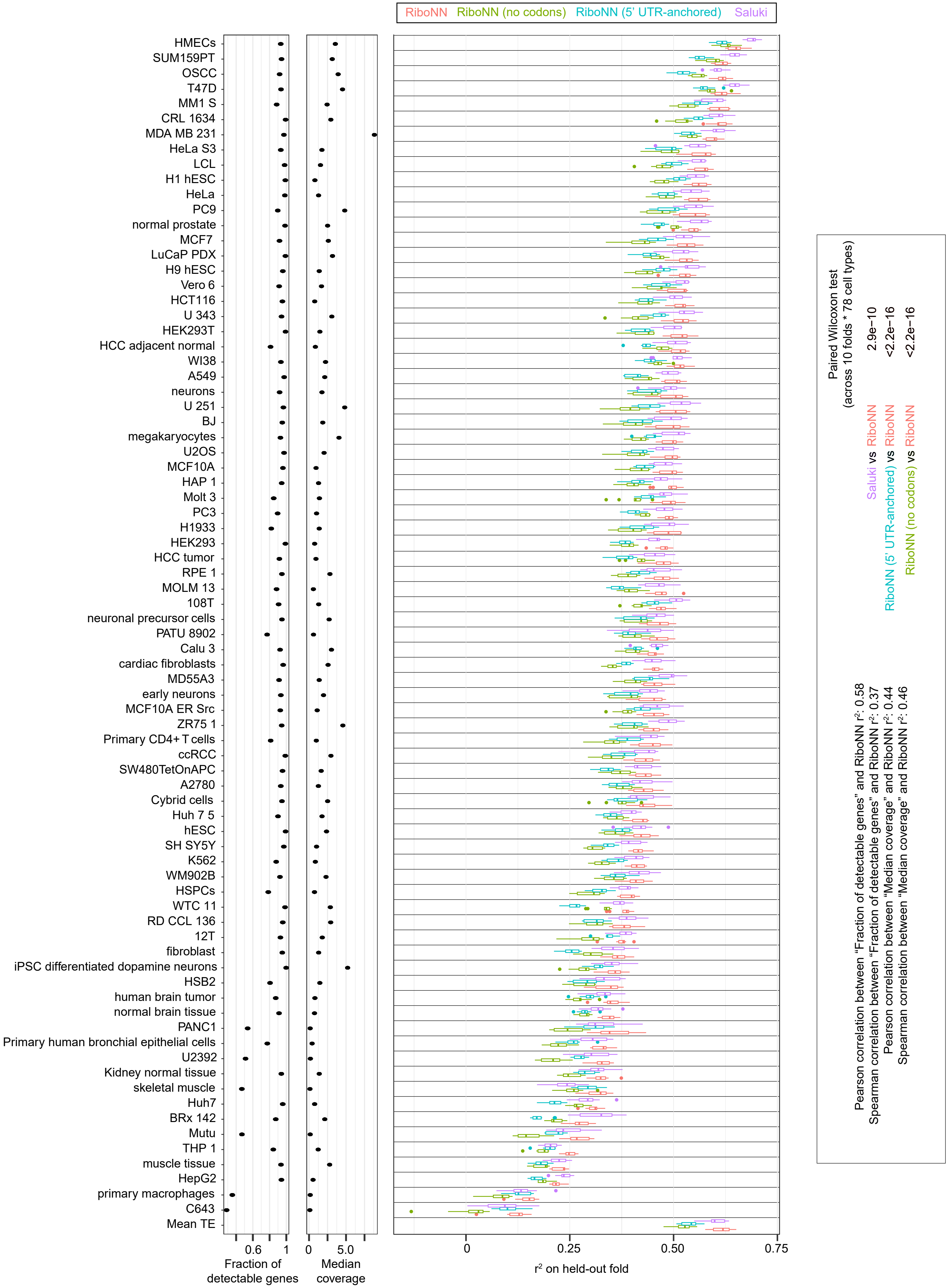
RiboNN
RiboNN (no codons)
RiboNN (5' UTR-anchored)
Saluki
HMECs
SUM159PT
OSCC
T47D
MM1 S
CRL 1634
MDA MB 231
HeLa S3
LCL
H1 hESC
HeLa
PC9
normal prostate
MCF7
LuCaP PDX
H9 hESC
Vero 6
HCT116
U 343
HEK293T
HCC adjacent normal
WI38
A549
neurons
U 251
BJ
megakaryocytes
U2OS
MCF10A
HAP 1
Molt 3
PC3
H1933
HEK293
HCC tumor
RPE 1
MOLM 13
108T
neuronal precursor cells
PATU 8902
Calu 3
cardiac fibroblasts
MD55A3
early neurons
MCF10A ER Src
ZR75 1
Primary CD4+ T cells
ccRCC
SW480TetOnAPC
A2780
Cybrid cells
Huh 7 5
hESC
SH SY5Y
K562
WM902B
HSPCs
WTC 11
RD CCL 136
12T
fibroblast
iPSC differentiated dopamine neurons
HSB2
human brain tumor
normal brain tissue
PANC1
Primary human bronchial epithelial cells
U2392
Kidney normal tissue
skeletal muscle
Huh7
BRx 142
Mutu
THP 1
muscle tissue
HepG2
primary macrophages
C643
Mean TE
0.6
1
Fraction of detectable genes
0
5.0
Median coverage
0
0.25
0.50
0.75
r² on held-out fold
Paired Wilcoxon test
(across 10 folds * 78 cell types)
Saluki vs RiboNN 2.9e−10
RiboNN (5' UTR-anchored) vs RiboNN <2.2e−16
RiboNN (no codons) vs RiboNN <2.2e−16
Pearson correlation between "Fraction of detectable genes" and RiboNN r²: 0.58
Spearman correlation between "Fraction of detectable genes" and RiboNN r²: 0.37
Pearson correlation between "Median coverage" and RiboNN r²: 0.44
Spearman correlation between "Median coverage" and RiboNN r²: 0.46

**Extended Data Fig. 3. Performance of deep learning models on all human cell types.** These panels mirror those shown in **Supplementary Fig. 4**, except show the performance of multitask deep learning models with one of four architectures: i) our final RiboNN architecture, ii) our RiboNN architecture, ablating the input channel recording codon positions, iii) our RiboNN architecture, anchoring all mRNAs at their 5′ end instead of at start codons, and iv) the Saluki architecture[2], removing the splice site input channel.

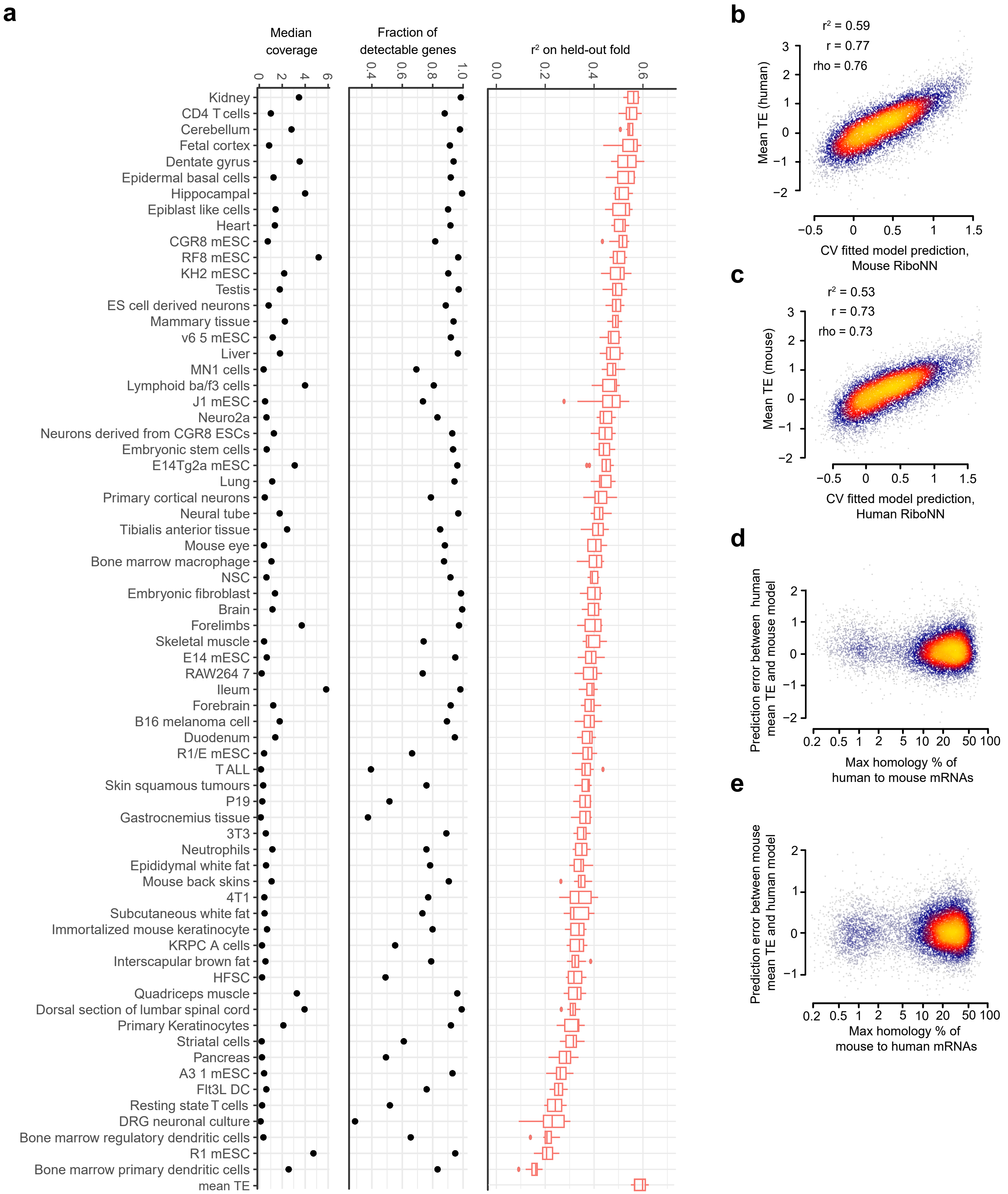

Pearson correlation between “Fraction of detectable genes” and RiboNN r²: 0.46
Spearman correlation between “Fraction of detectable genes” and RiboNN r²: 0.43
Pearson correlation between “Median coverage” and RiboNN r²: 0.22
Spearman correlation between “Median coverage” and RiboNN r²: 0.41

**Extended Data Fig. 4. Performance of RiboNN on mouse cell types. a,** These panels mirror those shown in **Supplementary Fig. 4**, except show the performance of our multitask RiboNN model on mouse cell types. **b-c,** Scatter plots showing the relationships between our mouse RiboNN predictions to the observed mean TEs for human mRNAs **(b)** as well as the relationships between our human RiboNN predictions to the observed mean TEs for mouse mRNAs **(c)**. Pearson (r) and Spearman (rho) correlation coefficients are also shown. **d-e,** Scatter plots showing the relationships between sequence homology, considering the interspecies pair of mRNAs with the maximum homology, and the residual prediction error between the TE from one species and TE predicted from the alternative species. This was shown for human **(d)** and mouse **(e)** mean TE data. "Max homology %" was computed as follows: i) all human–mouse mRNA pairs were locally aligned using the "pairwiseAlignment" function from the Biostrings (version 2.70.2) R package[3] ('match: 1, mismatch: −3, gap open: −2, and gap extend: −1'), and ii) for each mRNA, the final value was computed using the highest scoring alignment from the other species, calculating the maximum homology score divided by mRNA length.

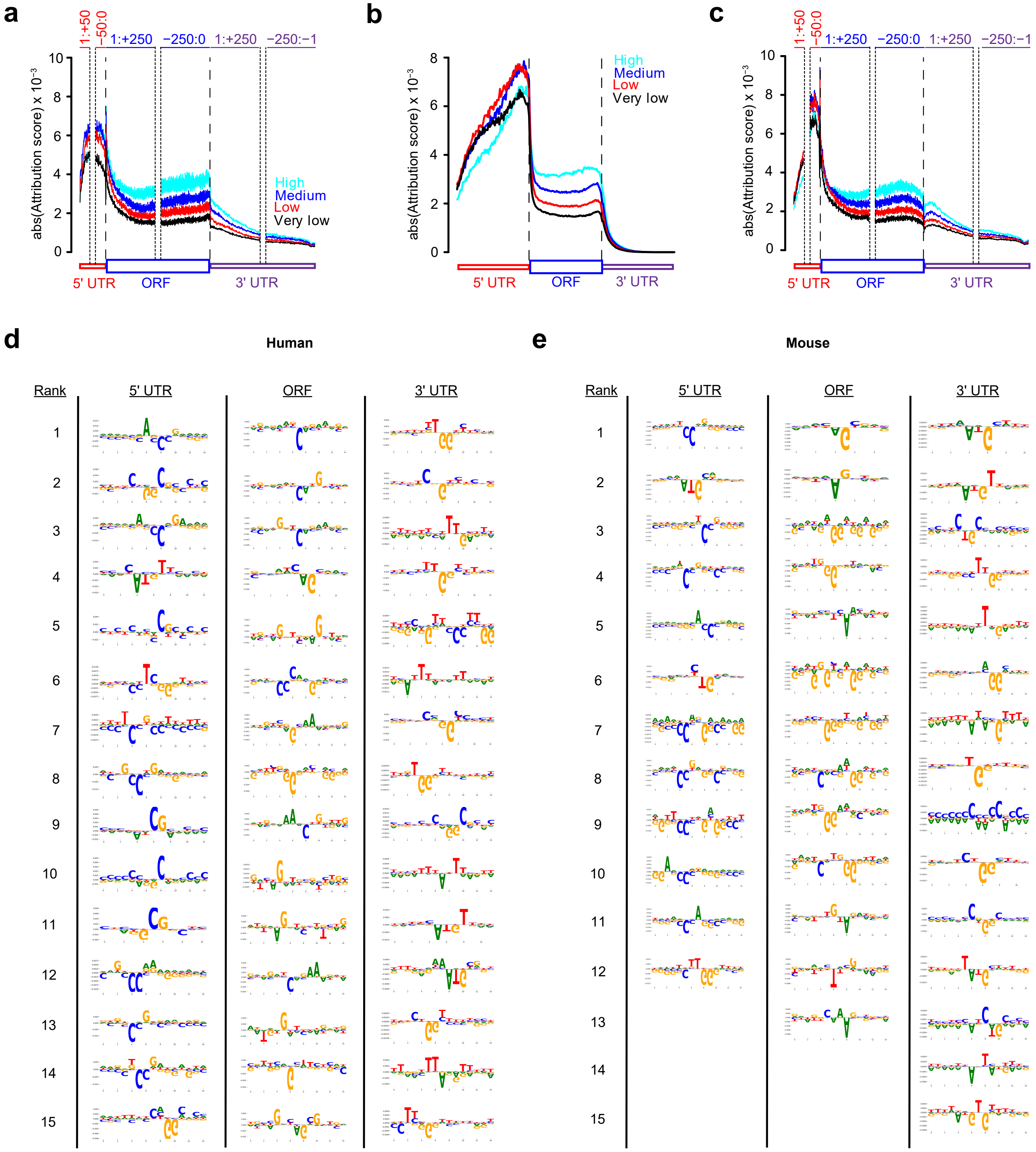
a
b
c
1:+50
-50:0
1:+250
−250:0
1:+250
−250:−1
abs(Attribution score) x 10⁻³
High
Medium
Low
Very low
5' UTR
ORF
3' UTR
d
Human
e
Mouse
Rank
5' UTR
ORF
3' UTR

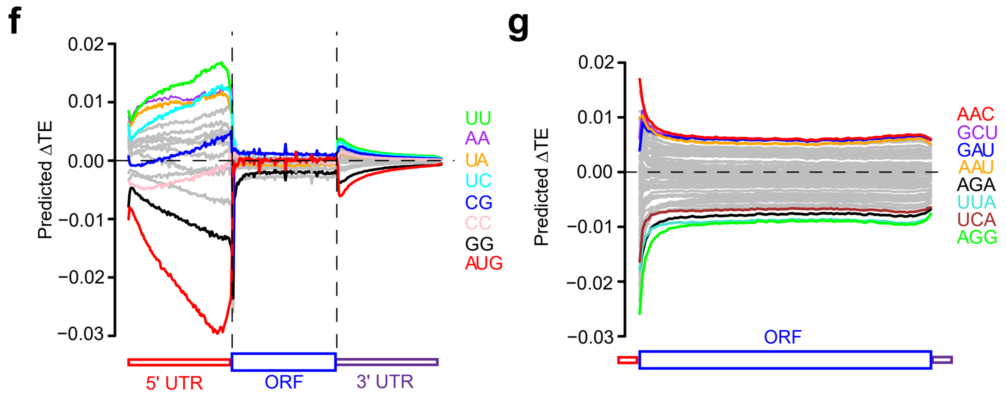

**Extended Data Fig. 5. Interpretation of human and mouse RiboNN models. a,** Attribution score plot for human RiboNN model focusing on specific regions along valid mRNAs (defined in **Methods**). The windows include the first 50 nt of the 5′ UTR, 50 nt upstream to 250 nt downstream of the start codon, 250 nt upstream and 250 nt downstream of the stop codon, and the last 250 nt of the 3′ UTR. The absolute values of attribution scores were averaged across all valid mRNAs, which were grouped into one of 4 equally sized bins according to their mean TE. **b,** Metagene plot for the absolute value of attribution scores derived from the mouse RiboNN model, averaged across all mRNAs, for percentiles along the 5′ UTR, CDS, and 3′ UTR. mRNAs were grouped into one of 4 equally sized bins according to their mean TE. **c,** This panel is the same as panel **a)**, except it reflects results from the mouse RiboNN model. **d-e,** Enriched motifs learned by human **(d)** and mouse **(e)** RiboNN models for each functional region of mRNA. Motifs are ranked by the number of seqlets[4] supporting each motif. **f,** Insertional analysis of 16 dinucleotides and the AUG motif for the mouse RiboNN model. Motifs were inserted into each of 100 equally spaced positional bins along the 5′ UTR, CDS, and 3′ UTRs of each mRNA. The average predicted change in TE for each bin is indicated and plotted along a metagene. **g,** This panel is the same as panel **(f)**, except it performs the analysis for 61 codons (excluding the 3 stop codons) inserted into the first reading frame along the length of the CDS.

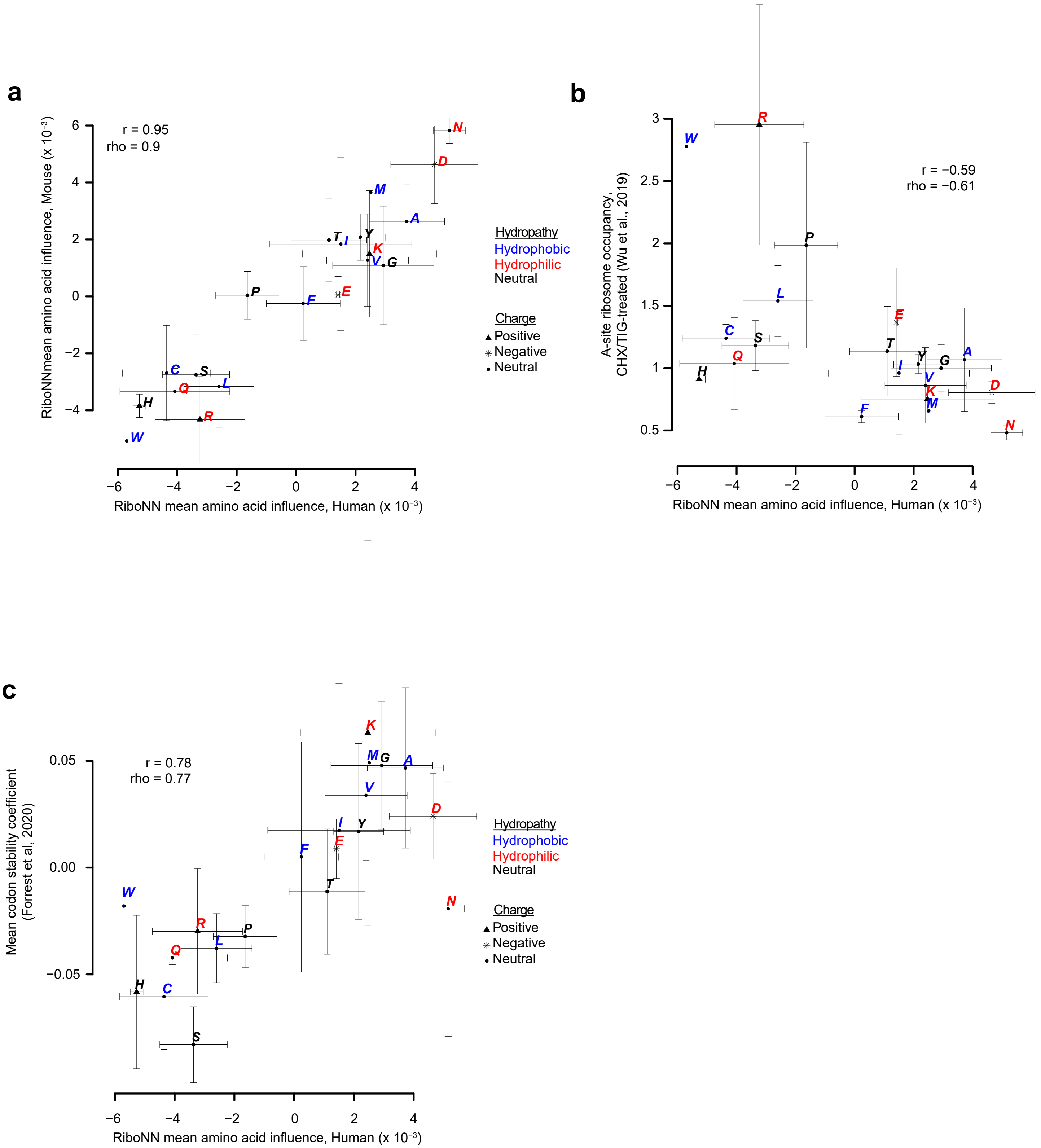

**Extended Data Fig. 6. Amino acid level-based correlation among codon influence scores. a-c,** Scatter plots showing the relationship between the amino acid level-based codon influence (*i.e.*, the predicted effect size of each inserted codon, averaged across all positional bins) from the human RiboNN model with that of the mouse model **(a)**, A-site ribosome occupancy scores[5] **(b)**, and mean codon stability coefficients[6] **(c)**. Pearson (r) and Spearman (rho) correlation coefficients are also shown. The properties of amino acids are labeled by different colors for hydrophobicity and by different shapes for charge. The error bar represents the standard error across codons encoding the same amino acid. To compute amino acid-level scores, we computed the mean score among codons encoding the same amino acid.

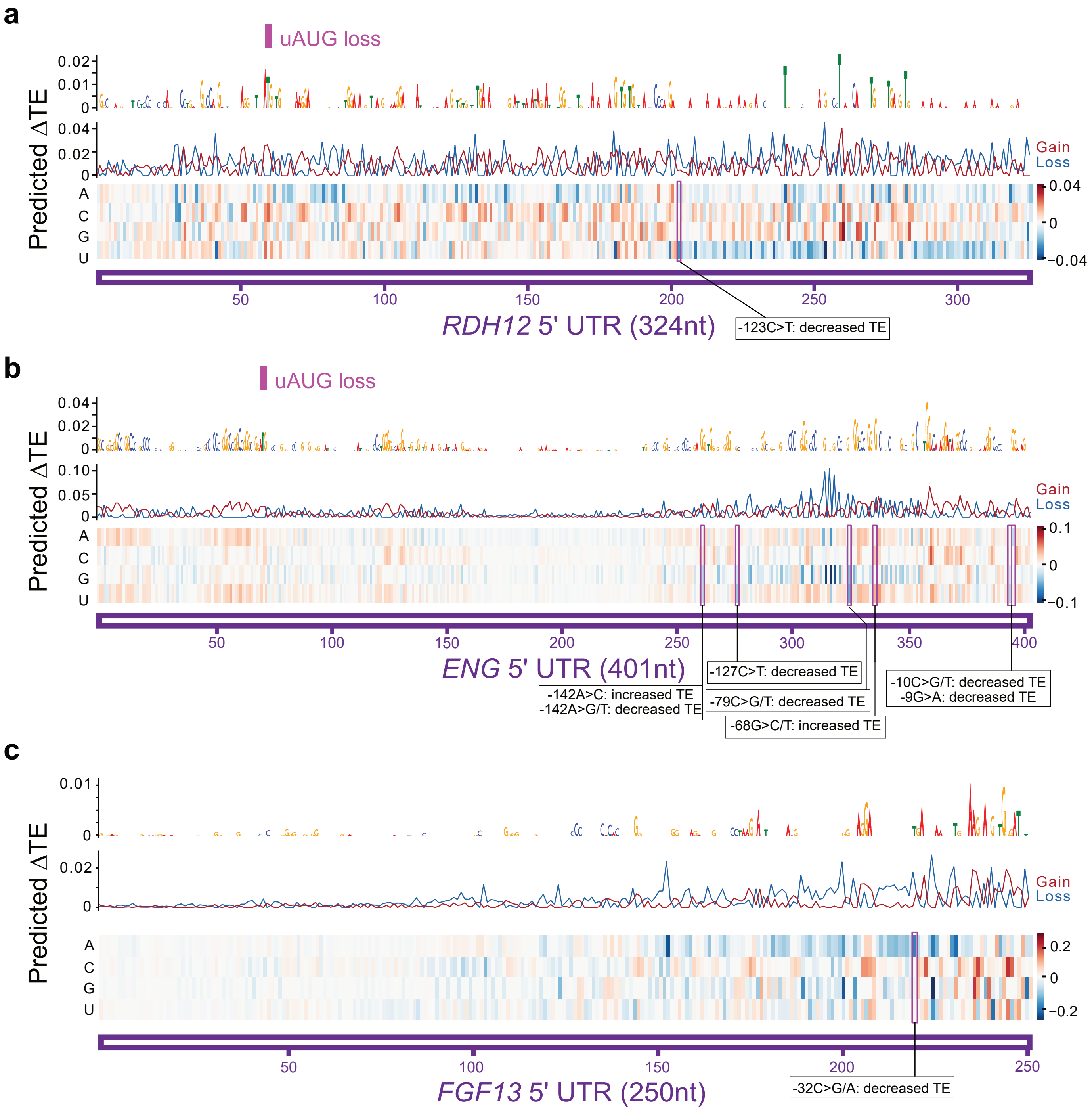
a
uAUG loss
Predicted ΔTE
Gain
Loss
A
C
G
U
RDH12 5' UTR (324nt)
-123C>T: decreased TE
b
uAUG loss
Predicted ΔTE
Gain
Loss
ENG 5' UTR (401nt)
-142A>C: increased TE
-142A>G/T: decreased TE
-127C>T: decreased TE
-79C>G/T: decreased TE
-68G>C/T: increased TE
-10C>G/T: decreased TE
-9G>A: decreased TE
c
Predicted ΔTE
Gain
Loss
FGF13 5' UTR (250nt)
-32C>G/A: decreased TE

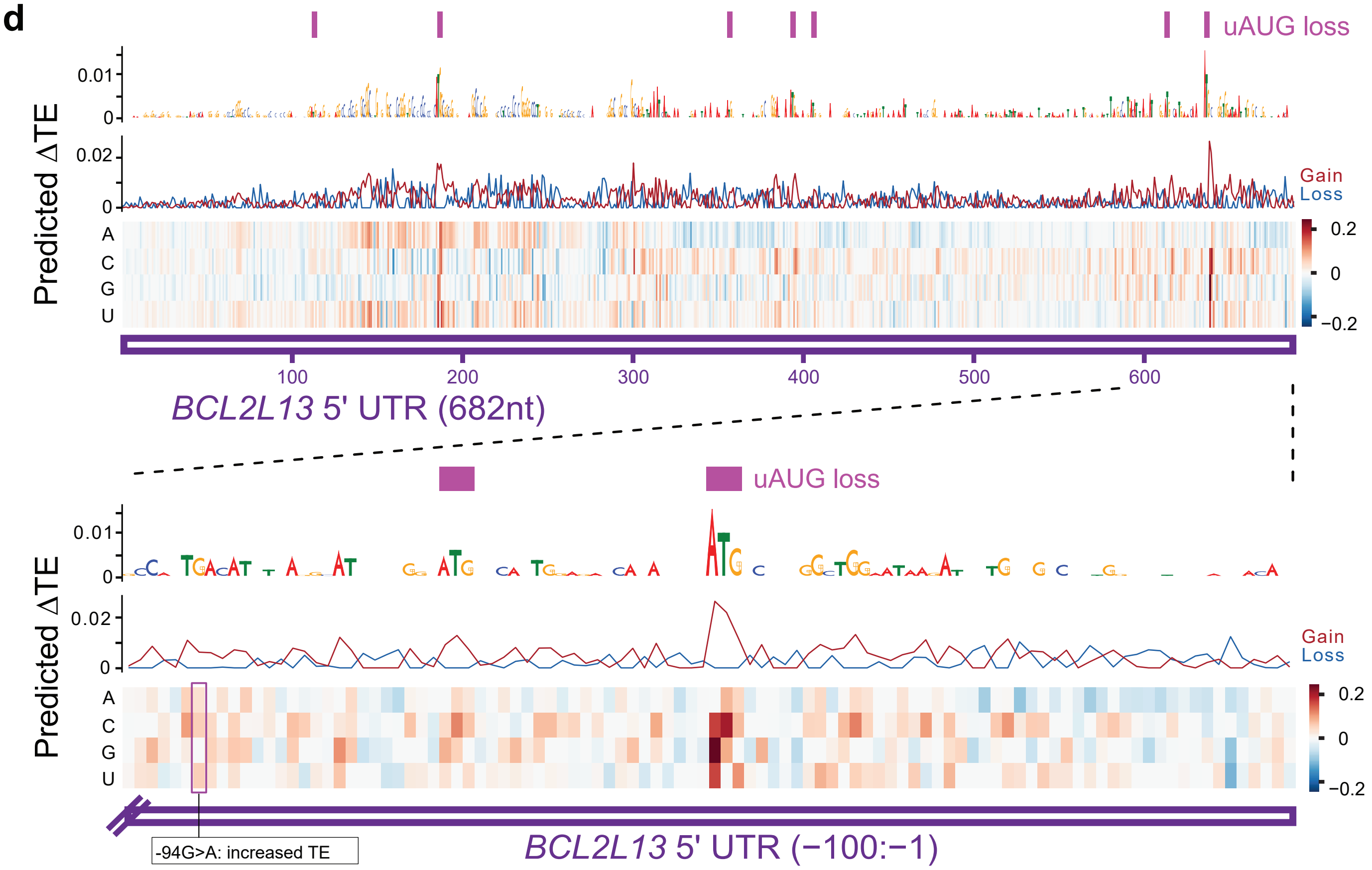

**Extended Data Fig. 7.** ***In silico*** **mutagenesis of disease-associated gene 5′ UTRs. a-d,** *In silico* mutagenesis of 5′ UTR regions of *RDH12* **(a)**, *ENG* **(b)**, *FGF13* **(c)**, and *BCL2L13* **(d)**. Positions of wild type uAUG are highlighted in purple at the top. The known disease-associated variants are boxed. Single point mutations resulting in predicted TE differences are shown alongside annotations reflecting the corresponding gain or loss of TE.

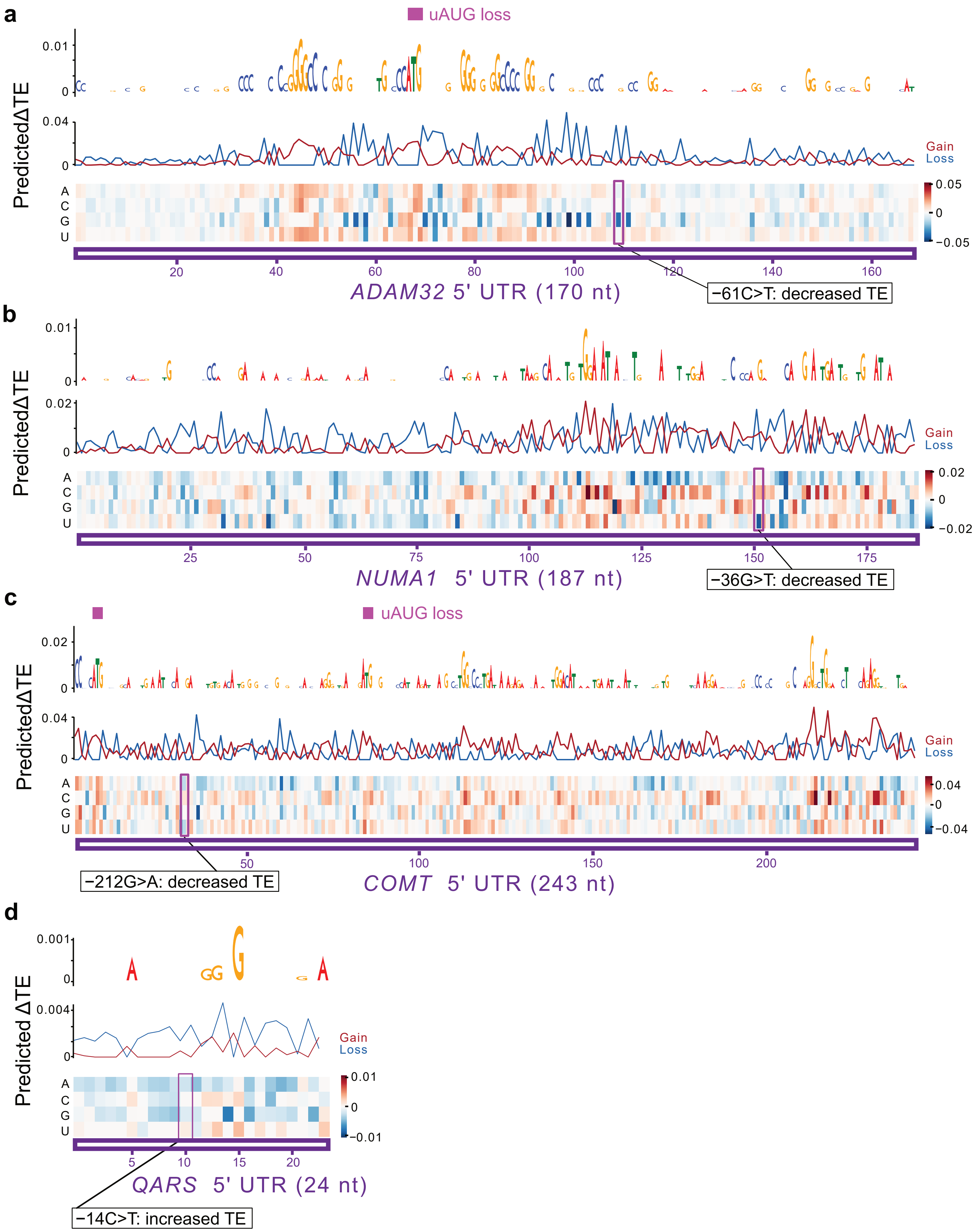

a
uAUG loss
Predicted∆TE
Gain
Loss
ADAM32 5' UTR (170 nt)
−61C>T: decreased TE
b
NUMA1 5' UTR (187 nt)
−36G>T: decreased TE
c
uAUG loss
COMT 5' UTR (243 nt)
−212G>A: decreased TE
d
QARS 5' UTR (24 nt)
−14C>T: increased TE

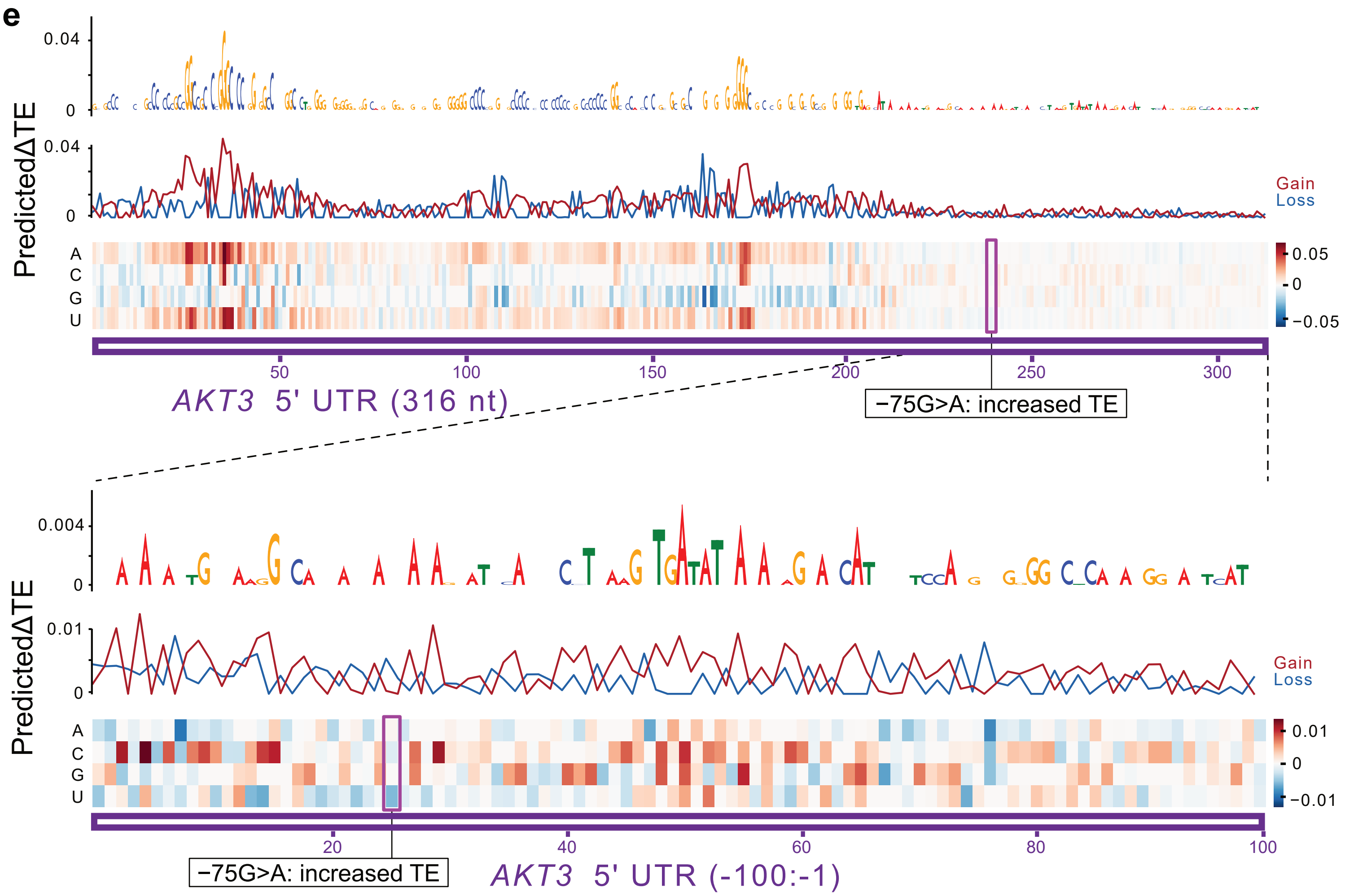

**Extended Data Fig. 8. *In silico* mutagenesis of cancer-associated gene 5′ UTRs. a-e,** *In silico* mutagenesis of 5′ UTR regions of *ADAM32* **(a)**, *NUMA1* **(b)**, *COMT* **(c)**, *QARS* **(d)**, and *AKT3* **(e)**. Positions of wild type uAUG are highlighted in purple at the top. The known cancer-associated variants are boxed. Single point mutations resulting in predicted TE differences are shown alongside annotations reflecting the corresponding gain or loss of TE.

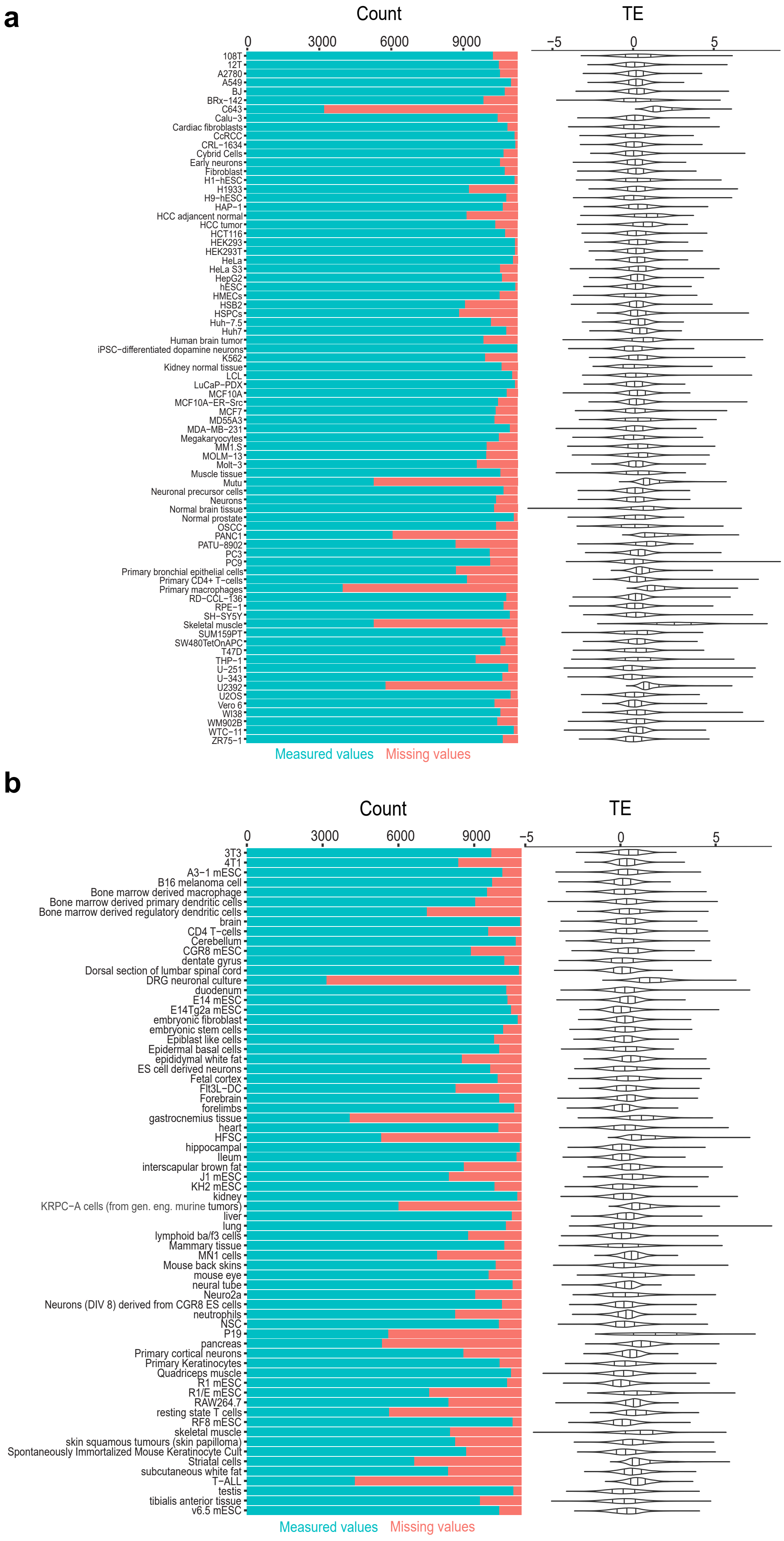
a
Count
TE
0
3000
6000
9000
−5
0
5
108T
12T
A2780
A549
BJ
BRx−142
C643
Calu−3
Cardiac fibroblasts
CcRCC
CRL−1634
Cybrid Cells
Early neurons
Fibroblast
H1−hESC
H1933
H9−hESC
HAP−1
HCC adjancent normal
HCC tumor
HCT116
HEK293
HEK293T
HeLa
HeLa S3
HepG2
hESC
HMECs
HSB2
HSPCs
Huh−7.5
Huh7
Human brain tumor
iPSC−differentiated dopamine neurons
K562
Kidney normal tissue
LCL
LuCaP−PDX
MCF10A
MCF10A−ER−Src
MCF7
MD55A3
MDA−MB−231
Megakaryocytes
MM1.S
MOLM−13
Molt−3
Muscle tissue
Mutu
Neuronal precursor cells
Neurons
Normal brain tissue
Normal prostate
OSCC
PANC1
PATU−8902
PC3
PC9
Primary bronchial epithelial cells
Primary CD4+ T−cells
Primary macrophages
RD−CCL−136
RPE−1
SH−SY5Y
Skeletal muscle
SUM159PT
SW480TetOnAPC
T47D
THP−1
U−251
U−343
U2392
U2OS
Vero 6
WI38
WM902B
WTC−11
ZR75−1
Measured values
Missing values
b
Count
TE
0
3000
6000
9000
−5
0
5
3T3
4T1
A3−1 mESC
B16 melanoma cell
Bone marrow derived macrophage
Bone marrow derived primary dendritic cells
Bone marrow derived regulatory dendritic cells
brain
CD4 T−cells
Cerebellum
CGR8 mESC
dentate gyrus
Dorsal section of lumbar spinal cord
DRG neuronal culture
duodenum
E14 mESC
E14Tg2a mESC
embryonic fibroblast
embryonic stem cells
Epiblast like cells
Epidermal basal cells
epididymal white fat
ES cell derived neurons
Fetal cortex
Flt3L−DC
Forebrain
forelimbs
gastrocnemius tissue
heart
HFSC
hippocampal
Ileum
interscapular brown fat
J1 mESC
KH2 mESC
kidney
KRPC−A cells (from gen. eng. murine tumors)
liver
lung
lymphoid ba/f3 cells
Mammary tissue
MN1 cells
Mouse back skins
mouse eye
neural tube
Neuro2a
Neurons (DIV 8) derived from CGR8 ES cells
neutrophils
NSC
P19
pancreas
Primary cortical neurons
Primary Keratinocytes
Quadriceps muscle
R1 mESC
R1/E mESC
RAW264.7
resting state T cells
RF8 mESC
skeletal muscle
skin squamous tumours (skin papilloma)
Spontaneously Immortalized Mouse Keratinocyte Cult
Striatal cells
subcutaneous white fat
T−ALL
testis
tibialis anterior tissue
v6.5 mESC
Measured values
Missing values

**Supplementary Fig. 1. Quality and distribution of mammalian TE measurements. a-b,** Stacked barplots (left panel) of the number of TE measurements and missing values out of 10,348 genes over 78 human cell types **(a)** and 10,870 genes over 68 mouse cell types **(b)**. Also shown is the distribution of available TE measurements over all cell types (right panel).

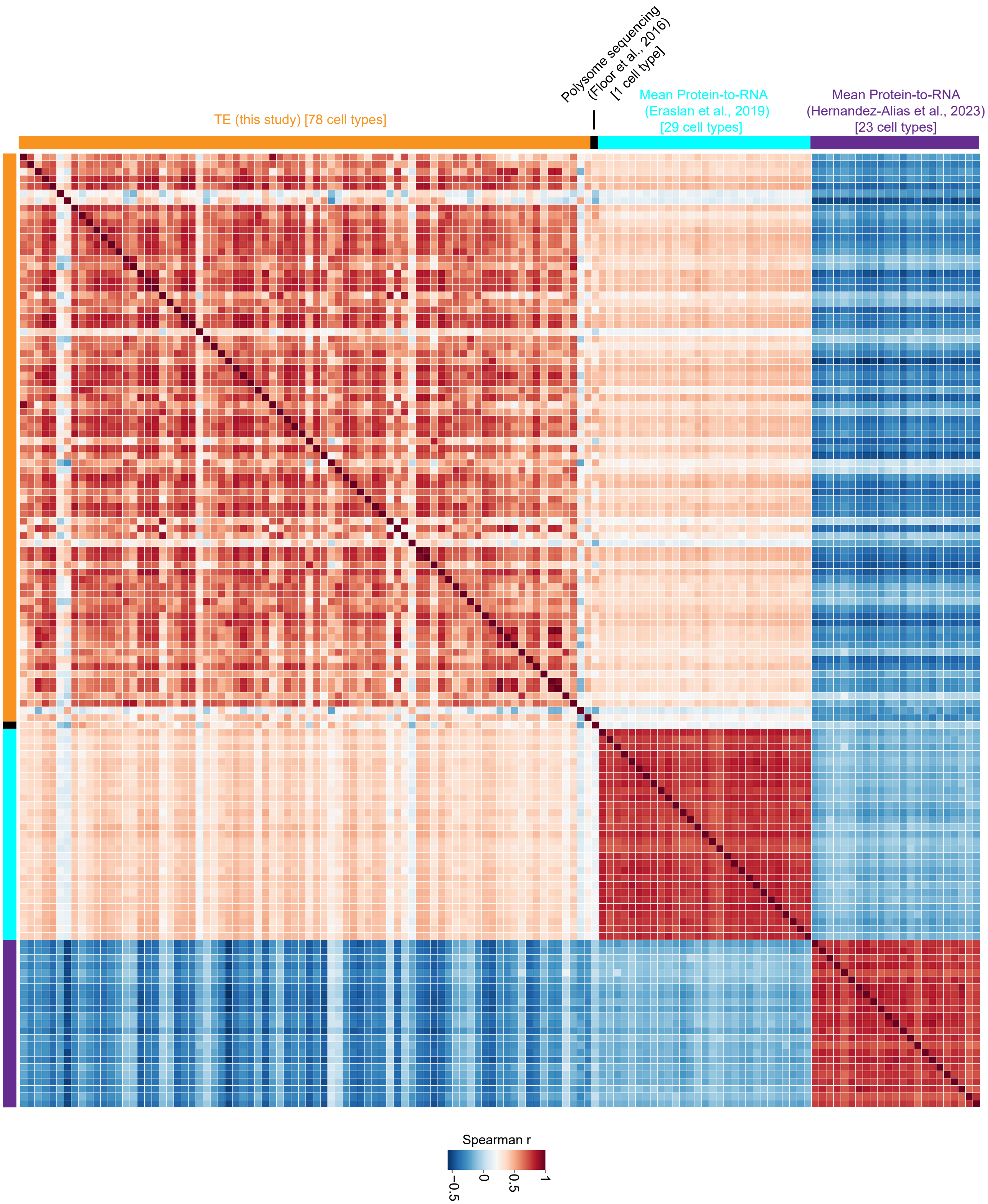

**Supplementary Fig. 2. Relationship among different measurements of translational output in diverse cell types.** This heatmap shows the correlations between TEs for mRNAs derived from this study and alternative measurements of translational output from prior studies[7–9] among numerous cell types. The colors are indicative of Spearman rho values.

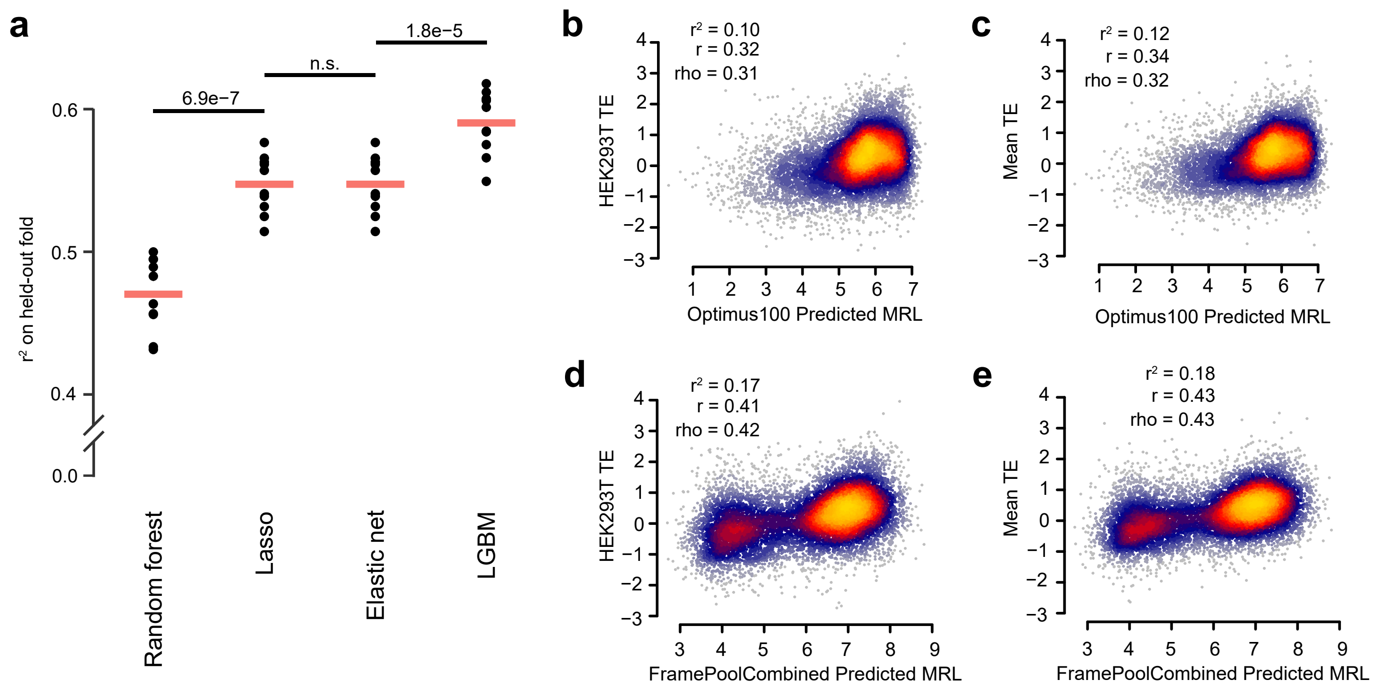

**Supplementary Fig. 3. Comparison of historical and classical machine learning methods. a,** Benchmarking of multiple ML approaches in prediction performance. Performance was measured as the $r^2$ on the held-out fold of ten CV folds on the human mean TE task. Models used the optimal feature set determined in **Fig. 2a**. Significance tests were determined by one-sided, paired t-tests adjusted for multiple hypothesis testing with a Bonferroni correction. **b-e,** Performance of Optimus[10] (**b**-**c**) and FramePool[11] (**d**-**e**) models in predicting HEK293T or mean TEs based upon 5′ UTR sequences.

a

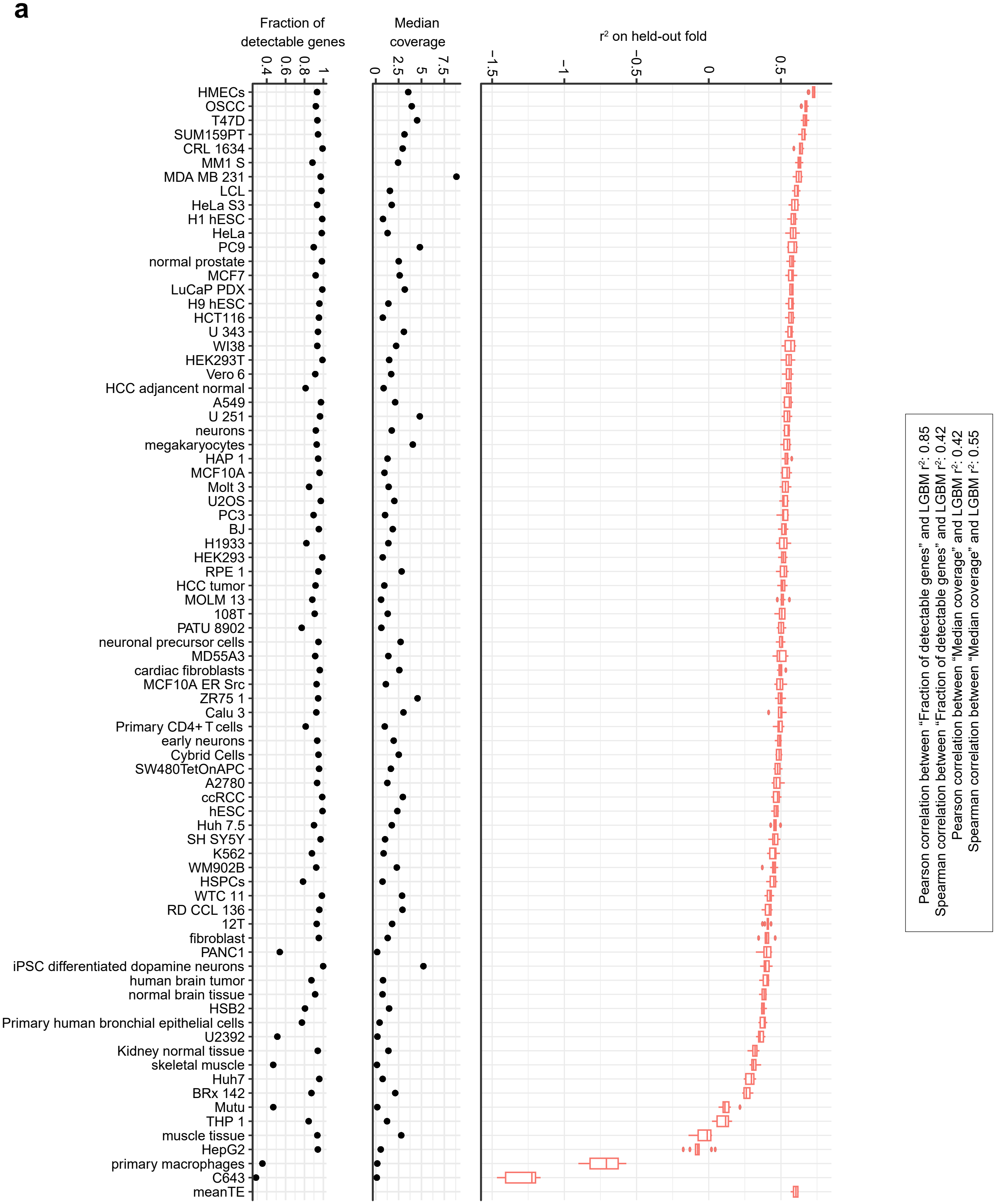

Fraction of detectable genes
Median coverage
r² on held-out fold
0.4 0.6 0.8 1
0 2.5 5 7.5
−1.5 −1 −0.5 0 0.5
HMECs
OSCC
T47D
SUM159PT
CRL 1634
MM1 S
MDA MB 231
LCL
HeLa S3
H1 hESC
HeLa
PC9
normal prostate
MCF7
LuCaP PDX
H9 hESC
HCT116
U 343
WI38
HEK293T
Vero 6
HCC adjancent normal
A549
U 251
neurons
megakaryocytes
HAP 1
MCF10A
Molt 3
U2OS
PC3
BJ
H1933
HEK293
RPE 1
HCC tumor
MOLM 13
108T
PATU 8902
neuronal precursor cells
MD55A3
cardiac fibroblasts
MCF10A ER Src
ZR75 1
Calu 3
Primary CD4+ T cells
early neurons
Cybrid Cells
SW480TetOnAPC
A2780
ccRCC
hESC
Huh 7.5
SH SY5Y
K562
WM902B
HSPCs
WTC 11
RD CCL 136
12T
fibroblast
PANC1
iPSC differentiated dopamine neurons
human brain tumor
normal brain tissue
HSB2
Primary human bronchial epithelial cells
U2392
Kidney normal tissue
skeletal muscle
Huh7
BRx 142
Mutu
THP 1
muscle tissue
HepG2
primary macrophages
C643
meanTE
Pearson correlation between “Fraction of detectable genes” and LGBM r²: 0.85
Spearman correlation between “Fraction of detectable genes” and LGBM r²: 0.42
Pearson correlation between “Median coverage” and LGBM r²: 0.42
Spearman correlation between “Median coverage” and LGBM r²: 0.55

b

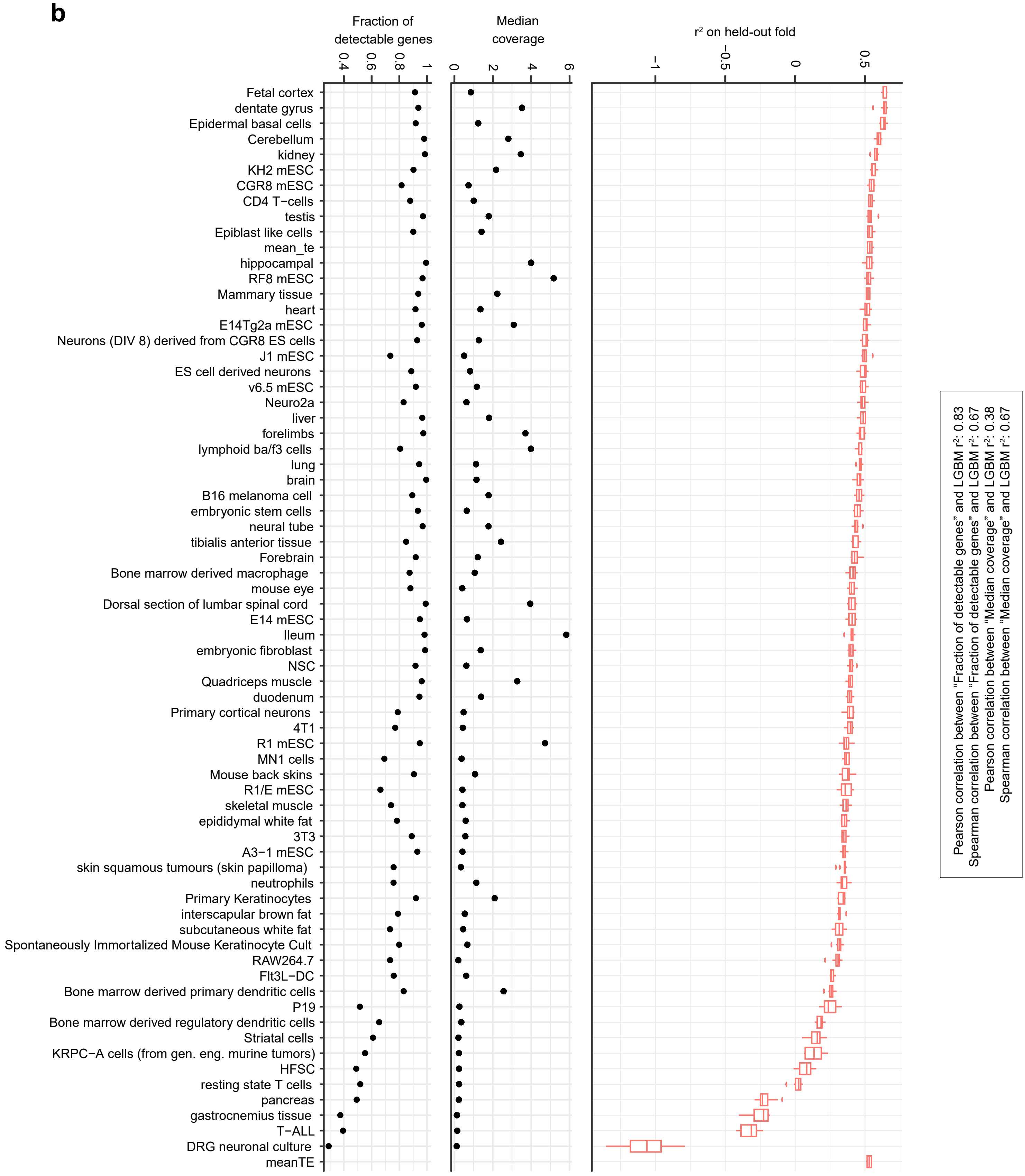

Fraction of detectable genes
0.4
0.6
0.8
1
Median coverage
0
2
4
6
r² on held-out fold
−1
−0.5
0
0.5
Fetal cortex
dentate gyrus
Epidermal basal cells
Cerebellum
kidney
KH2 mESC
CGR8 mESC
CD4 T−cells
testis
Epiblast like cells
mean_te
hippocampal
RF8 mESC
Mammary tissue
heart
E14Tg2a mESC
Neurons (DIV 8) derived from CGR8 ES cells
J1 mESC
ES cell derived neurons
v6.5 mESC
Neuro2a
liver
forelimbs
lymphoid ba/f3 cells
lung
brain
B16 melanoma cell
embryonic stem cells
neural tube
tibialis anterior tissue
Forebrain
Bone marrow derived macrophage
mouse eye
Dorsal section of lumbar spinal cord
E14 mESC
Ileum
embryonic fibroblast
NSC
Quadriceps muscle
duodenum
Primary cortical neurons
4T1
R1 mESC
MN1 cells
Mouse back skins
R1/E mESC
skeletal muscle
epididymal white fat
3T3
A3−1 mESC
skin squamous tumours (skin papilloma)
neutrophils
Primary Keratinocytes
interscapular brown fat
subcutaneous white fat
Spontaneously Immortalized Mouse Keratinocyte Cult
RAW264.7
Flt3L−DC
Bone marrow derived primary dendritic cells
P19
Bone marrow derived regulatory dendritic cells
Striatal cells
KRPC−A cells (from gen. eng. murine tumors)
HFSC
resting state T cells
pancreas
gastrocnemius tissue
T−ALL
DRG neuronal culture
meanTE
Pearson correlation between "Fraction of detectable genes" and LGBM r²: 0.83
Spearman correlation between "Fraction of detectable genes" and LGBM r²: 0.67
Pearson correlation between "Median coverage" and LGBM r²: 0.38
Spearman correlation between "Median coverage" and LGBM r²: 0.67

**Supplementary Fig. 4. Performance of LGBM models on all mammalian cell types. a-b,** Quality control of data for all human **(a)** and mouse **(b)** cell types (left and middle panels). Performance (right panel), measured as the $r^2$ on each of the ten held-out folds, between LGBM model predictions and observed TEs for each cell type. Models were trained using optimal feature set determined in **Fig. 2a**. Also shown is the correlation between quality control data and model performance across cell types (right box).

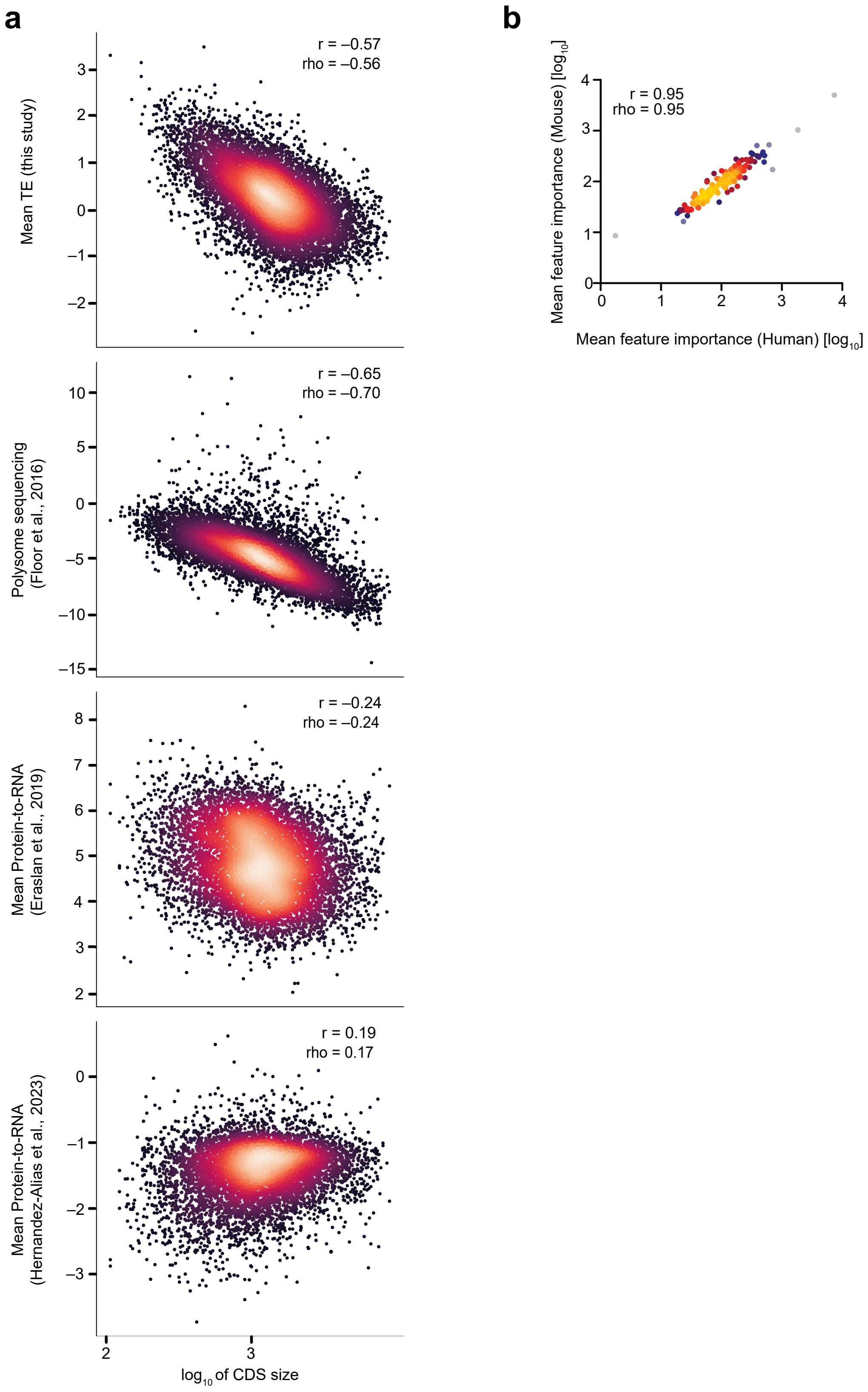

**Supplementary Fig. 5. Evaluation of LGBM features predictive of TE. a,** Scatter plots showing the relationships between ORF length and mean TE (from this study), polysome sequencing[7], and PTR data[8,9]. All datasets displayed a negative correlation to ORF length with one exception[9] which we previously found to be an outlier relative to all other measurements of translation rate (**Fig. 1c**). **b,** Scatter plot comparing mean feature importance of human and mouse LGBM models, as computed in **Methods**. Models used the optimal feature set determined in **Fig. 2a**.

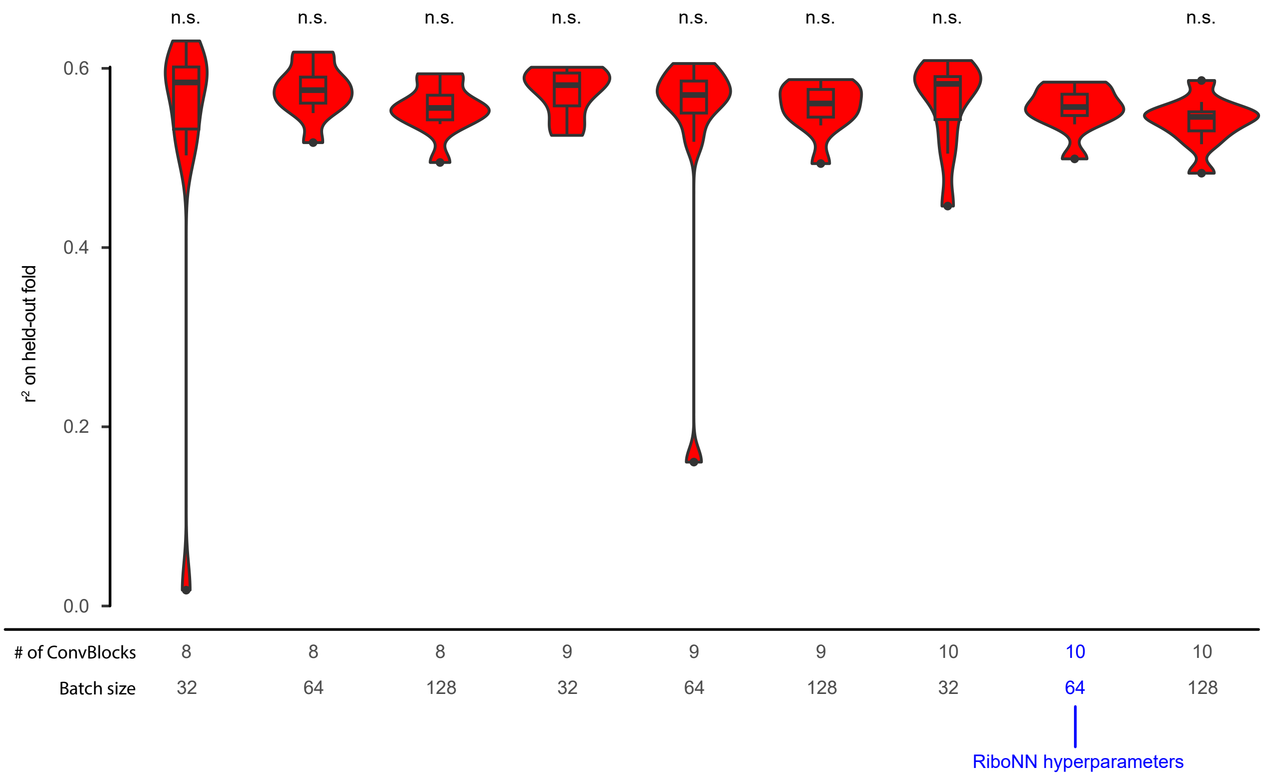

**Supplementary Fig. 6. Impact of hyperparameter choice on the performance of models that predict TE averaged across cell/tissue-types.** Violin plots with embedded boxplots showing the $r^2$ of models trained with different hyperparameter combinations (*e.g.*, varying the number of ConvBlocks and batch sizes) to predict mean TE. A paired Wilcoxon test, considering 10 held-out folds, was utilized to assess performance differences relative to RiboNN hyperparameters (shown in blue), with "n.s." indicating no significant difference.

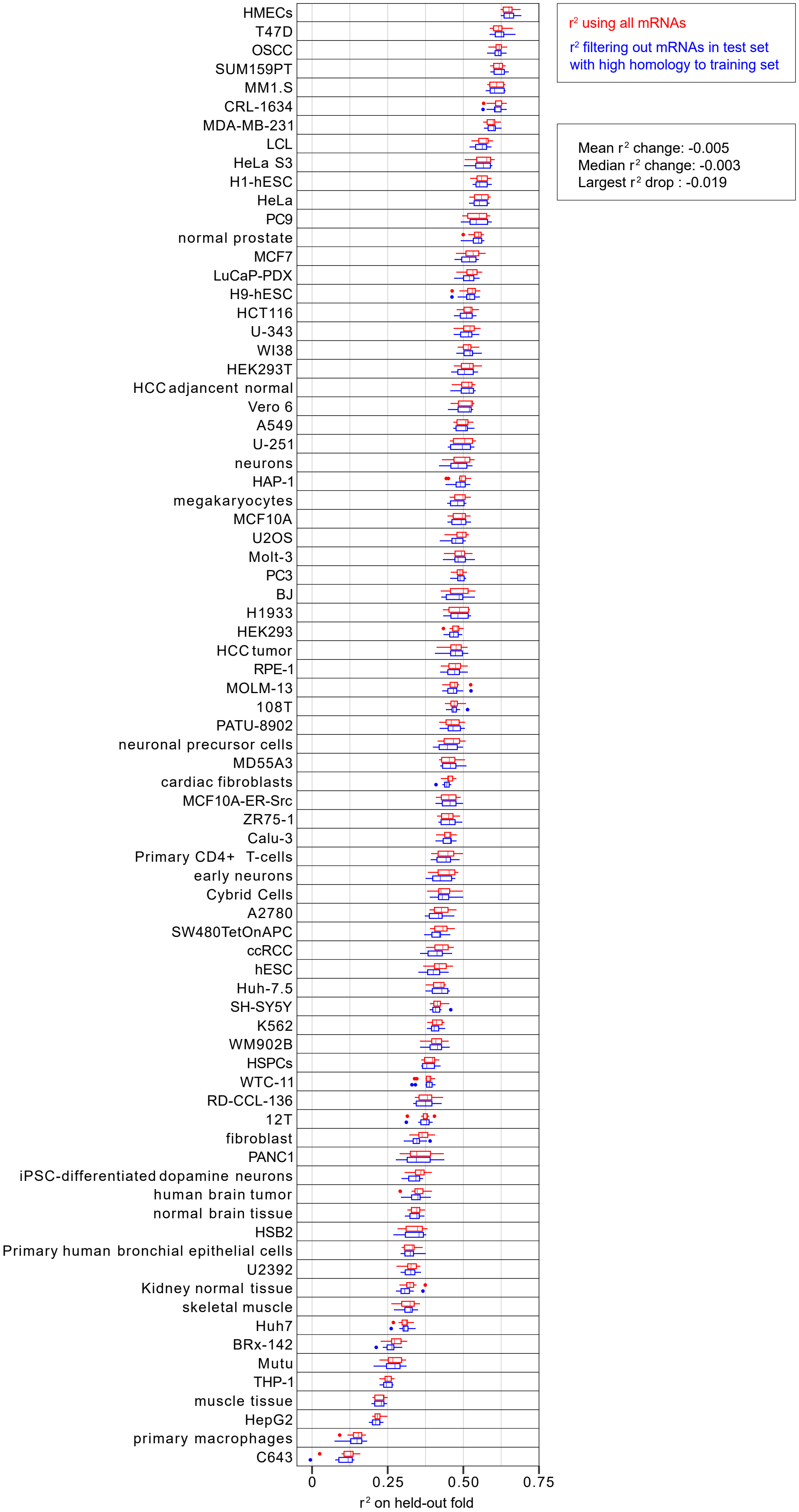
r2 using all mRNAs
r2 filtering out mRNAs in test set with high homology to training set
Mean r2 change: -0.005
Median r2 change: -0.003
Largest r2 drop : -0.019
HMECs
T47D
OSCC
SUM159PT
MM1.S
CRL-1634
MDA-MB-231
LCL
HeLa S3
H1-hESC
HeLa
PC9
normal prostate
MCF7
LuCaP-PDX
H9-hESC
HCT116
U-343
WI38
HEK293T
HCC adjancent normal
Vero 6
A549
U-251
neurons
HAP-1
megakaryocytes
MCF10A
U2OS
Molt-3
PC3
BJ
H1933
HEK293
HCC tumor
RPE-1
MOLM-13
108T
PATU-8902
neuronal precursor cells
MD55A3
cardiac fibroblasts
MCF10A-ER-Src
ZR75-1
Calu-3
Primary CD4+ T-cells
early neurons
Cybrid Cells
A2780
SW480TetOnAPC
ccRCC
hESC
Huh-7.5
SH-SY5Y
K562
WM902B
HSPCs
WTC-11
RD-CCL-136
12T
fibroblast
PANC1
iPSC-differentiated dopamine neurons
human brain tumor
normal brain tissue
HSB2
Primary human bronchial epithelial cells
U2392
Kidney normal tissue
skeletal muscle
Huh7
BRx-142
Mutu
THP-1
muscle tissue
HepG2
primary macrophages
C643
0
0.25
0.50
0.75
r2 on held-out fold

**Supplementary Fig. 7. Homology between mRNAs in the training and test sets does not strongly inflate prediction results.** Box plots comparing $r^2$ performance of RiboNN on original test folds (red) to test folds filtered to remove homologs (blue) over all human cell lines. Filtered test folds removed the subset of all mRNAs that shared ≥50% identity to an mRNA in the corresponding training folds. Percent nucleotide identity was computed as follows: i) each mRNA sequence in the test fold was aligned to all mRNA sequences in the training folds using the Biopython (v1.84)[12] PairwiseAligner module, implementing the Gotoh global alignment algorithm (parameters 'match=5, mismatch=−4, gap open=−10, and gap extend=−0.5'), and ii) for each mRNA, the percent nucleotide identity was computed using the highest scoring alignment, calculating the number of matched nucleotides divided by total alignment length.

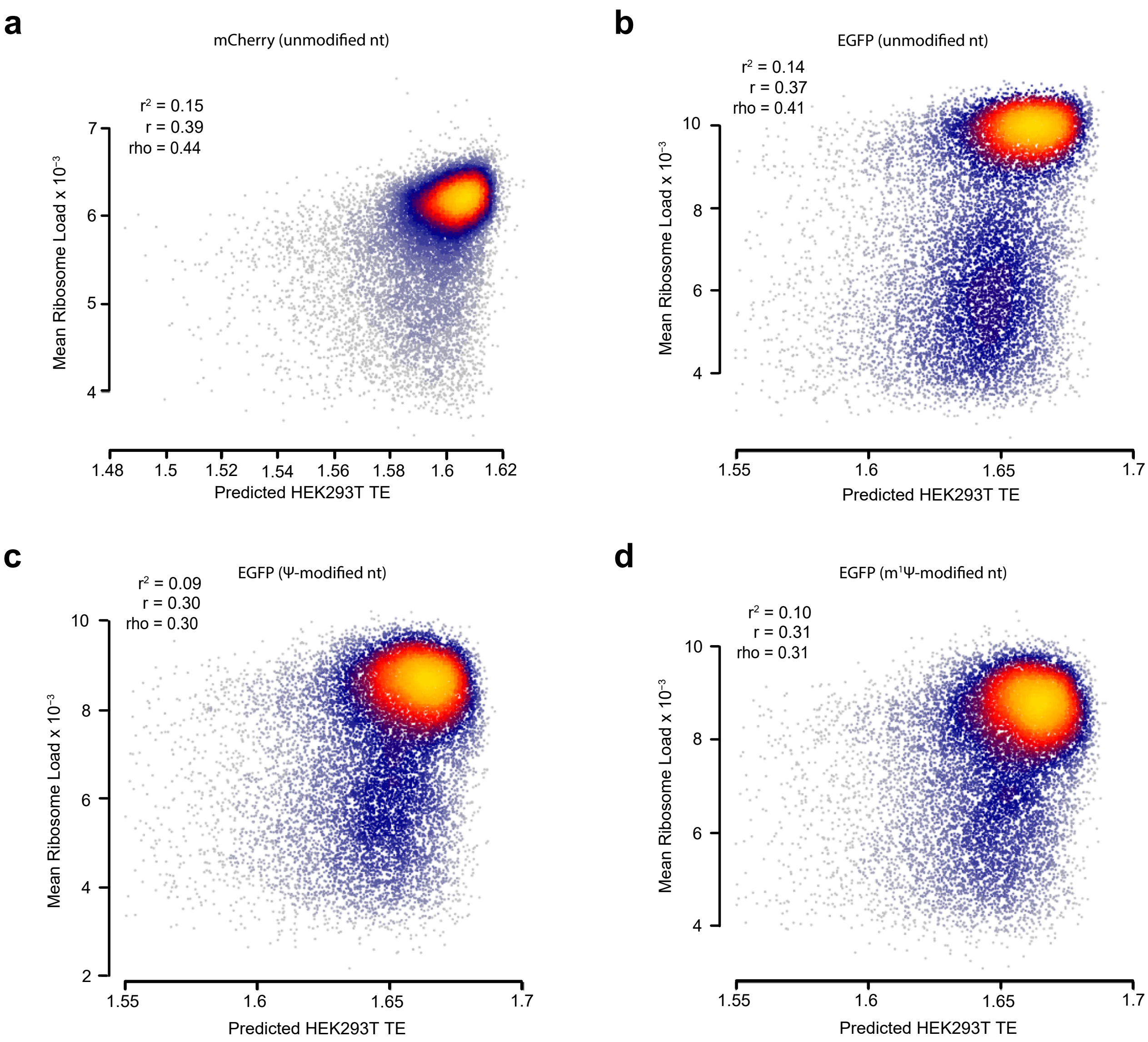

**Supplementary Fig. 9. Validation of human RiboNN using massively parallel reporter assay data. a-d,** Scatter plot showing the relationship of predicted TE from the HEK293T RiboNN model and experimentally measured mean ribosome load for reporter mRNAs[10] encoding mCherry **(a)**, EGFP **(b)**, Ψ-modified EGFP **(c)**, or m1Ψ-modified EGFP **(d).** Pearson (r) and Spearman (rho) correlation coefficients are also shown.

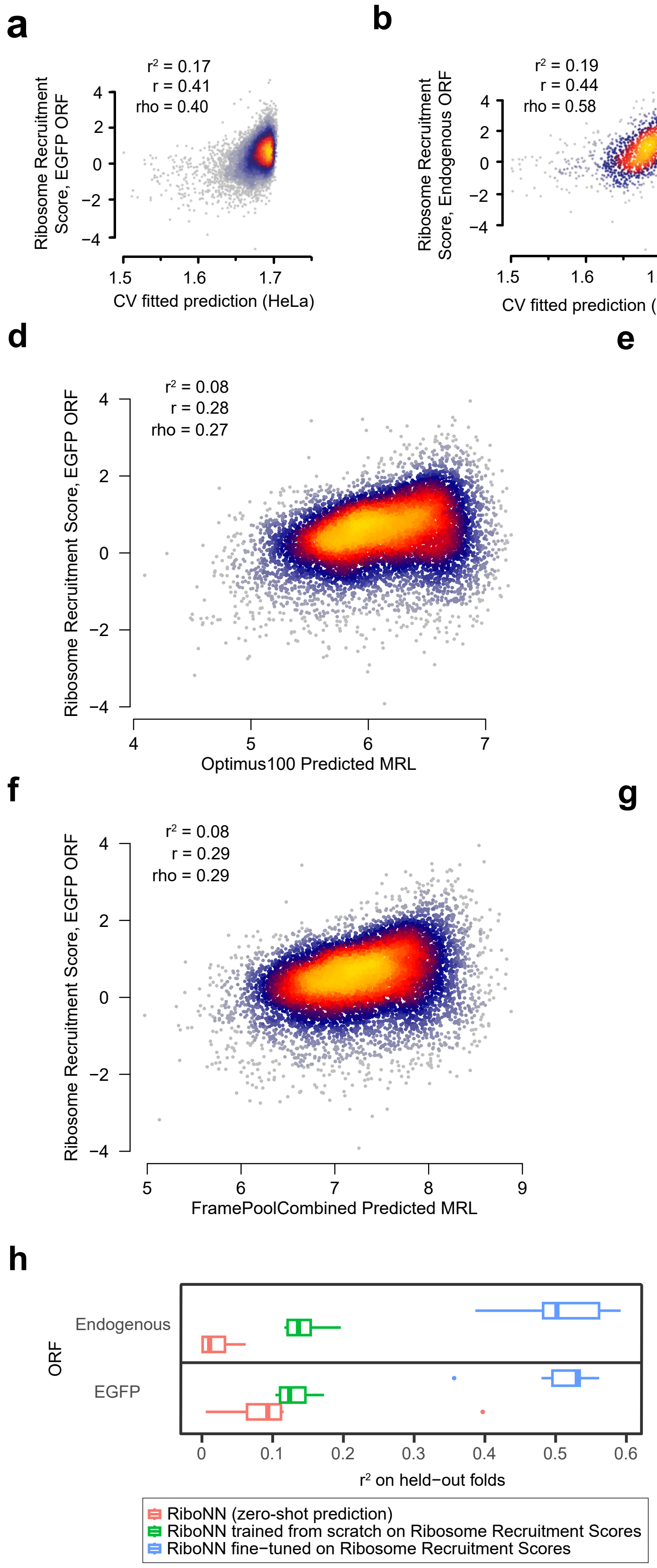

a
Ribosome Recruitment Score, EGFP ORF
r² = 0.17
r = 0.41
rho = 0.40
CV fitted prediction (HeLa)
b
Ribosome Recruitment Score, Endogenous ORF
r² = 0.19
r = 0.44
rho = 0.58
CV fitted prediction (HeLa)
c
Ribosome Recruitment Score, (Endogenous − EGFP ORF)
r² = 0.11
r = 0.33
rho = 0.39
Predicted ΔTE
d
Ribosome Recruitment Score, EGFP ORF
r² = 0.08
r = 0.28
rho = 0.27
Optimus100 Predicted MRL
e
Ribosome Recruitment Score, Endogenous ORF
r² = 0.03
r = 0.17
rho = 0.15
Optimus100 Predicted MRL
f
Ribosome Recruitment Score, EGFP ORF
r² = 0.08
r = 0.29
rho = 0.29
FramePoolCombined Predicted MRL
g
Ribosome Recruitment Score, Endogenous ORF
r² = 0.04
r = 0.19
rho = 0.19
FramePoolCombined Predicted MRL
h
ORF
Endogenous
EGFP
r² on held−out folds
RiboNN (zero-shot prediction)
RiboNN trained from scratch on Ribosome Recruitment Scores
RiboNN fine−tuned on Ribosome Recruitment Scores

**Supplementary Fig. 10. Benchmarking models to predict ribosome recruitment. a-c,** Performance of RiboNN in predicting the ribosomal recruitment score (*i.e.*, association of the 80S ribosomal subunit) to a panel of $m^1\Psi$-modified 5′ UTRs linked to EGFP (**a**), their corresponding endogenous ORFs (**b**), or the paired difference between the endogenous and EGFP ORF[16] (**c**). **d-g,** Performance of Optimus[10] (**d**-**e**) and FramePool[11] (**f**-**g**) models in predicting ribosome recruitment scores based upon 5′ UTR sequences linked to endogenous and EGFP ORFs. **h,** Performance of the pre-trained RiboNN model (zero-shot predictor), freshly trained RiboNN predictor, and fine-tuned RiboNN model on prediction of ribosome recruitment scores for 5′ UTRs linked to either endogenous ORFs or EGFP[16].

a

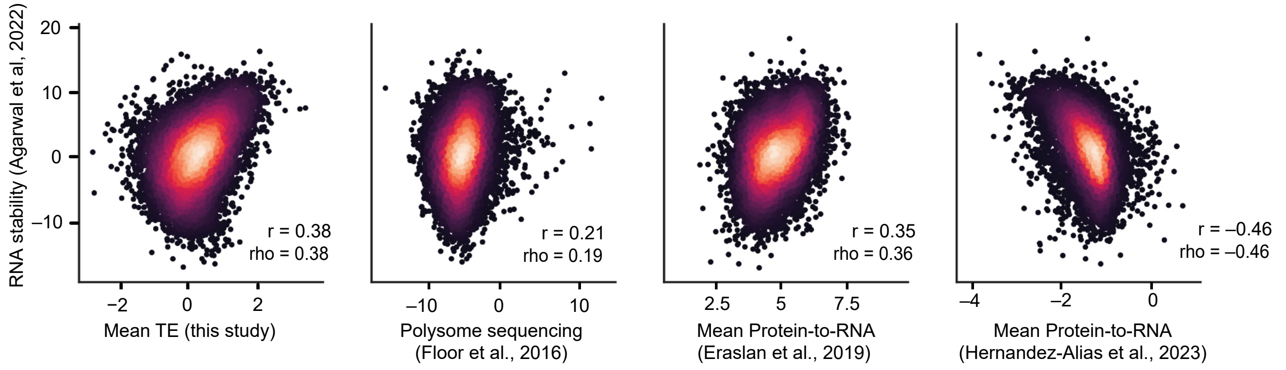

b

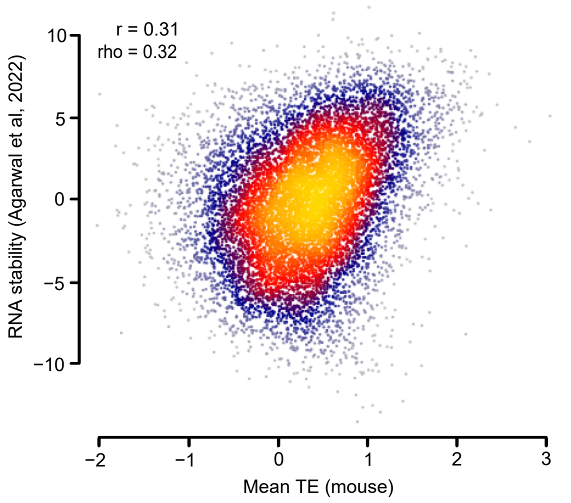

**Supplementary Fig. 11. Correlations between stability and translational measurements. a,** Scatter plots showing the relationships between human mRNA stability[2] and translational measurements including mean TE (from this study), polysome sequencing[7], and PTR data[8,9]. **b,** Scatter plot for the correlation between mouse mRNA stability[2] and TE. Pearson (r) and Spearman (rho) correlation coefficients are shown.
